# Channel-independent function of UNC-9/INX in spatial arrangement of GABAergic synapses in *C. elegans*

**DOI:** 10.1101/2022.05.16.492157

**Authors:** Ardalan Hendi, Longgang Niu, Andrew Snow, Richard Ikegami, Zhao-Wen Wang, Kota Mizumoto

## Abstract

Precise synaptic connection of neurons with their targets is essential for the proper functioning of the nervous system. A plethora of signaling pathways act in concert to mediate the precise spatial arrangement of synaptic connections. Here we show a novel role for a gap junction protein in controlling tiled synaptic arrangement in the GABAergic motor neurons in *C. elegans*, in which their axons and synapses overlap minimally with their neighboring neurons within the same class. We found that while EGL-20/Wnt controls axonal tiling, their presynaptic tiling is mediated by a gap junction protein UNC-9/Innexin, that is localized at the presynaptic tiling border between neighboring DD neurons. Strikingly, the gap junction channel activity of UNC-9 is dispensable for its function in controlling tiled presynaptic patterning. While gap junctions are crucial for the proper functioning of the nervous system as channels, our finding uncovered the novel channel-independent role of UNC-9 in synapse patterning.

## Introduction

Precise neuronal innervation and synaptic connections with their target cells are essential for proper functioning of the nervous system. During development, neurons communicate with their neighboring neurons to define their innervation pattern. Neuronal tiling is one type of inter-neuronal communications observed in many neuronal types, where neurons extend axons or dendrites in a non-overlapping manner with those from the neighboring neurons within the same class (Cameron and Rao, 2010; Grueber et al., 2002; Grueber and Sagasti, 2010; Grueber et al., 2003; Zipursky and Grueber, 2013). Distinct types of neuronal tiling and their regulators have been reported in many neuronal types across species. For example, in the visual system of *Drosophila*, L1 lamina neuron axons are arranged in columns in the medulla such that they only form synaptic connections within a single column in a non-redundant manner (Millard et al., 2007). Down syndrome cell adhesion molecule 2, DSCAM2, mediates axonal tiling of L1 lamina neuron axons through contact-dependent repulsive interactions between neighboring L1 neurons (Millard et al., 2007). Similarly, DSCAM serves as a homophilic repulsive signal to mediate self-avoidance and tiling in the mouse retinal amacrine cells (Fuerst et al., 2009; Fuerst et al., 2008). R7 photoreceptor neurons in the *Drosophila* visual system tile with neighboring R7 neurons through the TGFβ/activin signaling pathway (Ting et al., 2007). In *Drosophila*, the dendrites of neighboring class IV dendritic arborization neurons extend their dendrites in a non-overlapping manner with their neighboring neurons within the same class through Furry, Hippo and Tricornered (Emoto et al., 2004; Emoto et al., 2006). However, due to the technical limitations in labeling two neighboring neurons within the same class, our knowledge of genetic mechanisms that underlie neuronal tiling is still limited.

Tiling also occurs at the level of synapses. In *C. elegans*, dorsal-anterior DA motor neurons form *en passant* cholinergic chemical synapses onto the dorsal body wall muscles in a way that each presynaptic domain from a single DA neuron does not overlap with those from the neighboring DA neurons (White et al., 1986). Previously, we showed that Semaphorin and Plexin-dependent inter-axonal interaction defines the presynaptic tiling between two posterior DA neurons by locally inhibiting synapse formation (Mizumoto and Shen, 2013a).

Neurons use various conserved signaling and cell adhesion molecules for precise spatial arrangement of chemical synapses (Sanes and Yamagata, 2009; Yogev and Shen, 2014). Mutations in these genes lead to the formation of aberrant number of chemical synapses, which may underlie various neurodevelopmental and psychiatric disorders including autism spectrum disorder (ASD), schizophrenia and bipolar disorder (Guilmatre et al., 2009; Mitchell, 2011; Sudhof, 2008; Tabuchi et al., 2007; Tang et al., 2014; Wen et al., 2014). Several works showed that neurons use inhibitory cues to locally restrict synapse formation. For example, Sema3F and its receptors, Neurophilin-2 and PlexinA3, locally inhibit synapse formation in the proximal dendritic regions of cortical layer V pyramidal neurons (Tran et al., 2009). In *Drosophila*. Wnt4 secreted from the M13 muscles controls specificity of neuromuscular junctions by locally inhibiting synapse formation (Inaki et al., 2007). In *C. elegans,* two Wnts, LIN-44 and EGL-20 determine the topographic presynaptic arrangement of the DA-class of cholinergic motor neurons (Klassen and Shen, 2007; Mizumoto and Shen, 2013b). UNC-6/Netrin and its receptor UNC-5/DCC are required to inhibit presynaptic assembly in the DA9 dendrites (Poon et al., 2008).

In addition to chemical synapses, neurons also form electrical synapses through gap junction channels that mediate electrical coupling between the cells. Gap junctions consist of tetra-membrane spanning proteins, connexins (Cx) in mammals and innexins (INX) in invertebrates (Hall, 2017). Cx and INX monomers assemble into hexamers or octamers on neighboring cells that dock together to form gap junctions, through which neurons exchange small molecules and ions (Sanchez et al., 2019). We will hereafter refer to chemical synapses as synapses and electrical synapses as gap junctions, for simplicity. In addition, mammals have another family of gap junction proteins called pannexin (PANX), which share sequence similarity with INX. Unlike Cx and INX that form gap junction channels, PANXs only form hemichannels that mediate the exchange of small molecules and ions between the cytoplasm and extracellular space (Deng et al., 2020; Michalski et al., 2020). While gap junction proteins function primarily as channels, growing evidence supports channel-independent roles as cytoskeletal regulators. For example, human Cx43 and Drosophila INX2/3/4 control B lymphocyte and border cell migration, respectively, independent of their channel activities (Falk et al., 2014; Machtaler et al., 2011; Miao et al., 2020). Mammalian Cx43 and Cx26 mediate glial migration through a channel-independent adhesive role (Elias et al., 2007). The channel-independent roles of gap junction proteins in the nervous system are not well known. Interestingly, alterations in gap junction function and activity are associated with synaptopathies manifested by abnormal chemical synapse numbers and functions (Lapato and Tiwari-Woodruff, 2018; Swayne and Bennett, 2016). For example, increased Cx43 expression is observed in the prefrontal cortex of the post-mortem brain tissues of ASD patients (Fatemi et al., 2008). A PANX1 mutation is found in patients with intellectual disabilities (Shao et al., 2016). Upregulation of Cx43 and PANX1 is also associated with Alzheimer’s disease (Giaume et al., 2019). Functional and structural interactions between synapses and gap junctions have been observed in many aspects of neurodevelopment and function, yet the molecular mechanisms are largely unknown (Pereda, 2014).

Previous electron microscopy (EM) reconstruction of the *C. elegans* nervous system revealed axonal and dendritic tiling of the dorsal and ventral D-type (DDs and VDs) GABAergic motor neurons (White et al., 1986). Here we developed a system to stably label two neighboring DD motor neurons (DD5 and DD6) with fluorescent markers. Using this system, we show a unique combinatory regulation of axonal and presynaptic tiling by EGL-20/Wnt and UNC-9/INX. In *egl-20* mutants, axonal tiling between DD5 and DD6 was severely disrupted, while their presynaptic tiling was largely unaffected. We found that loss of *unc-9*, which encodes an INX gap junction protein, causes ectopic synapse formation in the distal axon of DD5, that resulting in the disrupted tiled presynaptic patterning in the *egl-20* mutant background. Strikingly, mutant UNC-9 proteins that form either putative constitutively closed or open gap junction channels could still rescue the presynaptic patterning defect of *unc-9* mutants, indicating that UNC-9’s gap junction channel activity is dispensable for its function in controlling presynaptic tiling. Our results reveal a novel channel-independent role for a gap junction protein in controlling synapse patterning. As UNC-9 gap junctions are formed at the DD5 and DD6 presynaptic tiling border, we propose that UNC-9 serves as a positional cue to define presynaptic domains.

## Results

### Tiled axonal, dendritic and presynaptic patterning of the DD5 and DD6 neurons

Cell bodies of six DD-class of GABAergic motor neurons (DD1 to DD6) reside in the ventral nerve cord (**Figure 1A**) (White et al., 1986). Each DD neuron extends a longer dendrite anteriorly and a shorter dendrite posteriorly, where they form postsynaptic dendritic spines. From the anterior dendrite, each DD neuron sends a commissure dorsally where it bifurcates to extend axons both anteriorly and posteriorly within the dorsal nerve cord. DD neurons form *en passant* synapses along their axons onto the dorsal body wall muscles and the VD motor neurons (White et al., 1986). Previous EM reconstruction has shown that the axons and dendrites of neighboring DD motor neurons have minimal overlap, where they form gap junction channels (White et al., 1986). As a result of tiled axonal and dendritic patterning, the domains of presynaptic and postsynaptic sites from each DD neuron do not overlap with those of the neighboring DD neurons, thereby achieving a tiled synaptic pattern.

**Figure 1:**
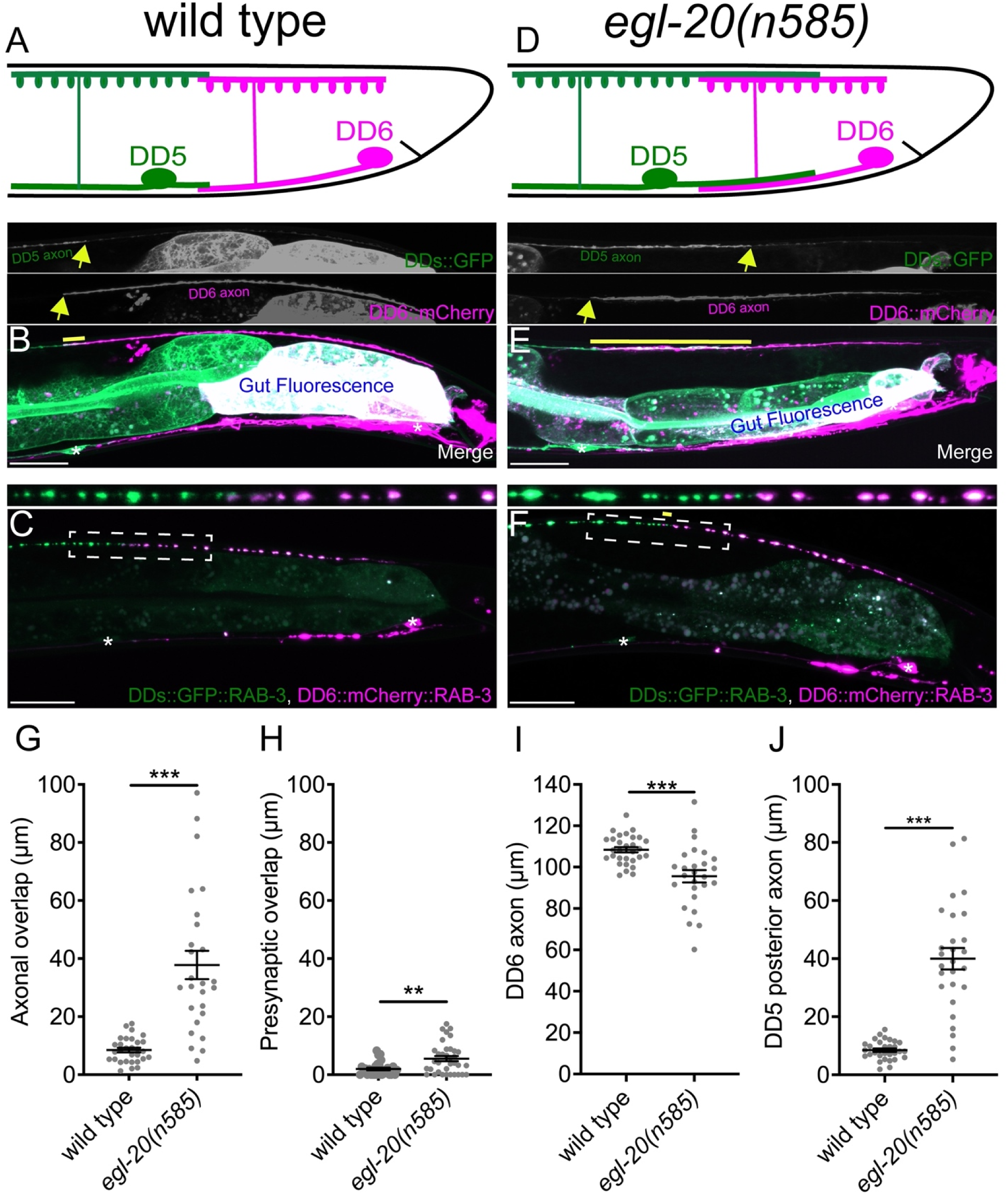
*egl-20/wnt* is required for axonal tiling between DD5 and DD6 neurons. (A) Schematic of axonal, dendritic and presynaptic tiling between DD5 and DD6 of wild type. (B) Representative image of axonal tiling in wild type animals. Yellow line represents region of axonal overlap between DD5 and DD6. Arrow indicates the end of DD5 posterior axon (top panel) and the end of DD6 anterior axon (middle panel). (C) Representative image of presynaptic tiling in wild type animals. The magnified straightened image of the presynaptic tiling border, represented by dotted box, is shown above. (D) Schematic of axonal and dendritic overlap between DD5 and DD6 of *egl-20(n585)* mutants. (E) Representative image of axonal tiling in *egl-20(n585)* mutant animals. Yellow line represents region of axonal overlap between DD5 and DD6. Arrow indicates the end of DD5 posterior axon (top panel) and the end of DD6 anterior axon (middle panel). (F) Representative image of presynaptic tiling in the *egl-20(n585)* mutants. Yellow line represents region of presynaptic overlap between DD5 and DD6 The magnified straightened image of the presynaptic tiling border, represented by dotted box, is shown above. Asterisks: DD5 and DD6 cell bodies. Scale bar: 20µm. (G) Quantification of axonal overlap between DD5 and DD6. (H) Quantification of presynaptic overlap between DD5 and DD6. (I) Quantification of DD6 axonal length. (J) Quantification of DD5 posterior axonal length. See Supplemental Figure 1-1A for the definition of the DD5 posterior axon. Each dot represents a single animal. Black bars indicate mean ± SEM. **p<0.01; ***p<0.001.

To visualize neurites of the two neighboring DD neurons in live animals, we created a transgenic marker stain in which all DD neurons express the membrane-associated GFP (GFP::CAAX) under the DD neuron-specific promoter (P*flp-13*), and the membrane-associated mCherry (mCherry::CAAX) under the DD6 neuron-specific promoter (P*plx-2*) (See Materials and Methods) (**Figure 1B**). The *flp-13* promoter activity in DD6 is often substantially dimmer than the rest of DD neurons (**Figure 1B**, personal communications with Michael Francis). This results in an increased color contrast between DD5, which expresses only GFP, and DD6, which expresses both mCherry and GFP. Using this marker strain, we observed minimal overlaps between the axons and the dendrites of DD5 and DD6 (**Figures 1A, 1B and 1G, Supplemental Figure 1-1**), which confirmed the previous electron microscopy data (White et al., 1986). To visualize presynaptic tiling of DD5 and DD6, we generated another transgenic marker strain, in which all DD neurons express GFP::RAB-3 presynaptic vesicle marker under the *flp-13* promoter, and DD6 expresses mCherry::RAB-3 under the *plx-2* promoter. Consistent with the tiled axonal projection pattern of DD5 and DD6, their presynaptic patterning also exhibited tiled innervation (**Figure 1A, 1C and 1H**).

Together, we showed for the first time, that DD5 and DD6 have tiled axons, dendrites, and presynaptic patterning in live animals, which are in agreement with the previous EM reconstruction data (White et al., 1986).

### EGL-20/Wnt inhibits the outgrowth of DD5 posterior axon and dendrite

We next asked what controls axonal and dendritic tiling between DD5 and DD6. Previous studies showed that Wnt signaling acts as a negative regulator of neurite outgrowth, including D-type motor neurons in *C. elegans* (Maro et al., 2009; Onishi et al., 2014; Zou, 2004). Among the five Wnt genes (*lin-44, egl-20, cwn-1, cwn-2, mom-2*) in *C. elegans*, *egl-20* is expressed in the cells around preanal ganglions (Whangbo and Kenyon, 1999), which are located near the DD5 and DD6 axonal and dendritic tiling border. In the loss-of-function mutants of *egl-20(n585)*, which carries a missense mutation in one of the conserved cysteine residues, we observed overextension of the DD5 posterior axon (**Figure 1D, 1E, 1J**), which resulted in significant overlaps between DD5 and DD6 axons (**Figures 1G**). Additionally, we also observed the overextention of the DD5 posterior dendrites in *egl-20(n585)* mutants (**Supplemental figure 1-1**). The overall structure of the DD6 neuron, including the length of DD6 axon is largely unaffected (**Figure 1I**), suggesting that EGL-20 specifically inhibits the outgrowth of the posterior axon and dendrite of DD5. Expression of *egl-20* from its endogenous promoter rescued the axonal overlap between DD5 and DD6 neurons (**Supplemental figure 1-2**). Similarly, the posterior dendrite of DD5 is also overextended in the *egl-20(n585)* mutants (**Figure 1D, 1E, Supplemental Figure 1-1**). We could not quantify the dendritic tiling defect due to the variable expression of mCherry::CAAX in the DD6 dendrite.

### Axonal and presynaptic tiling are controlled by different mechanisms

Since DD5 and DD6 axons overlap in the *egl-20(n585)* mutants, we asked whether their presynaptic patterns are also compromised. If the presynaptic tiling between DD5 and DD6 is simply a consequence of axonal tiling, the synapses will form throughout the DD5 axon in the *egl-20(n585)* mutants, resulting in significant overlap between the synaptic domains of DD5 and DD6. Surprisingly, despite the significant axonal overlap between DD5 and DD6, we observed little defect in the presynaptic tiling pattern (**Figure 1F and 1H**). While the degree of overlap between DD5 and DD5 presynaptic domains, which was defined as the distance between the most posterior DD5 presynaptic site and the most anterior DD6 presynaptic site, was significantly larger in the *egl-20(n585)* mutant compared with wild type, the small degree of presynaptic tiling defect did not reflect the large axonal overlap between DD5 and DD6 (**Figures 1G and 1H**). This observation strongly suggests that presynaptic tiling is not a consequence of axonal tiling, but rather there are additional mechanisms to tile their presynaptic domains even in the absence of axonal tiling.

As DD5 posterior dendrite also overextends in the *egl-20(n585)* mutants, we next examined the postsynaptic dendritic spine patterning of DD5 using a transgenic strain that expresses ACR-12::GFP under the *flp-13* promoter (Philbrook et al., 2018) (kind gift from M. Francis). ACR-12 is specifically localized at the postsynaptic sites of the GABAergic motor neurons dendritic spines (Barbagallo et al., 2017). Due to the dim expression from the *flp-13* promoter, ACR-12::GFP in DD6 is invisible in most animals, which allowed us to visualize the dendritic spines on the posterior dendrite of DD5. In wild type, the postsynaptic domain within the DD5 posterior dendrite, which we defined as the distance between DD5 cell body to the most posterior ACR-12::GFP punctum, is comparable to the length of DD5 posterior dendrite (**Supplemental figure 1-1**). This result indicates that the postsynaptic dendritic spines are formed throughout the length of DD5 posterior dendrite. In the *egl-20(n585)* mutant animals, the postsynaptic domain length in the DD5 posterior dendrite is significantly longer than in wild type, and roughly matched to the length of the DD5 posterior dendrite (**Supplemental figure 1-1J)**. While we could not examine the dendritic spine patterning of DD6 due to the weak expression of ACR-12::mCherry under the *plx-2* promoter, our results suggest that the position of DD5 postsynaptic dendritic spine is determined by the length of DD5 dendrite.

### UNC-9/INX is localized at the presynaptic tiling border between DD5 and DD6 axons

Previous electron microscopy studies have shown that gap junctions are formed in the region of minimal axonal overlap between DD neurons, where presynaptic tiling is established (White et al., 1976, 1986). These gap junctions are believed to play crucial roles in electrical coupling between DD neurons during sinusoidal locomotion (Kawano et al., 2011). DD neurons express several INX genes, including *unc-9* and *unc-7* (Altun et al., 2009). We first examined the subcellular localization of gap junctions between DD neurons by labelling the endogenous UNC-9 using the split-GFP based Native and Tissue-Specific Fluorescence (NATF) method (He et al., 2019). Briefly, we inserted seven tandem repeats of last β-strand of GFP (*7×gfp11*) at the C-terminus of endogenous locus of *unc-9* using CRISPR/Cas9 genome editing. The *unc-9(miz81; unc-9::7×gfp11)* animals exhibit uncoordinated locomotion pattern, probably because UNC-9::7×GFP11 is expected to form constitutively-open gap junction channels (see below). We then expressed the remaining part of GFP (GFP1-10) specifically in DD neurons using the *flp-13* promoter to reconstitute the fluorescent UNC-9::7×GFP exclusively in the DD neurons. In wild type animals, UNC-9::7×GFP was localized at the tip of anterior DD6 axon (**Figure 2A**). In the *egl-20(n585)* mutants, in which DD5 axon overextends to the DD6 axonal region, UNC-9::7×GFP was localized at the tip of DD6 axon similar to wild type (**Figure 2B**). This result suggests that axonal tiling defect in the *egl-20* mutants does not affect the position of UNC-9 gap junctions formed between DD5 and DD6 axons.

**Figure 2:**
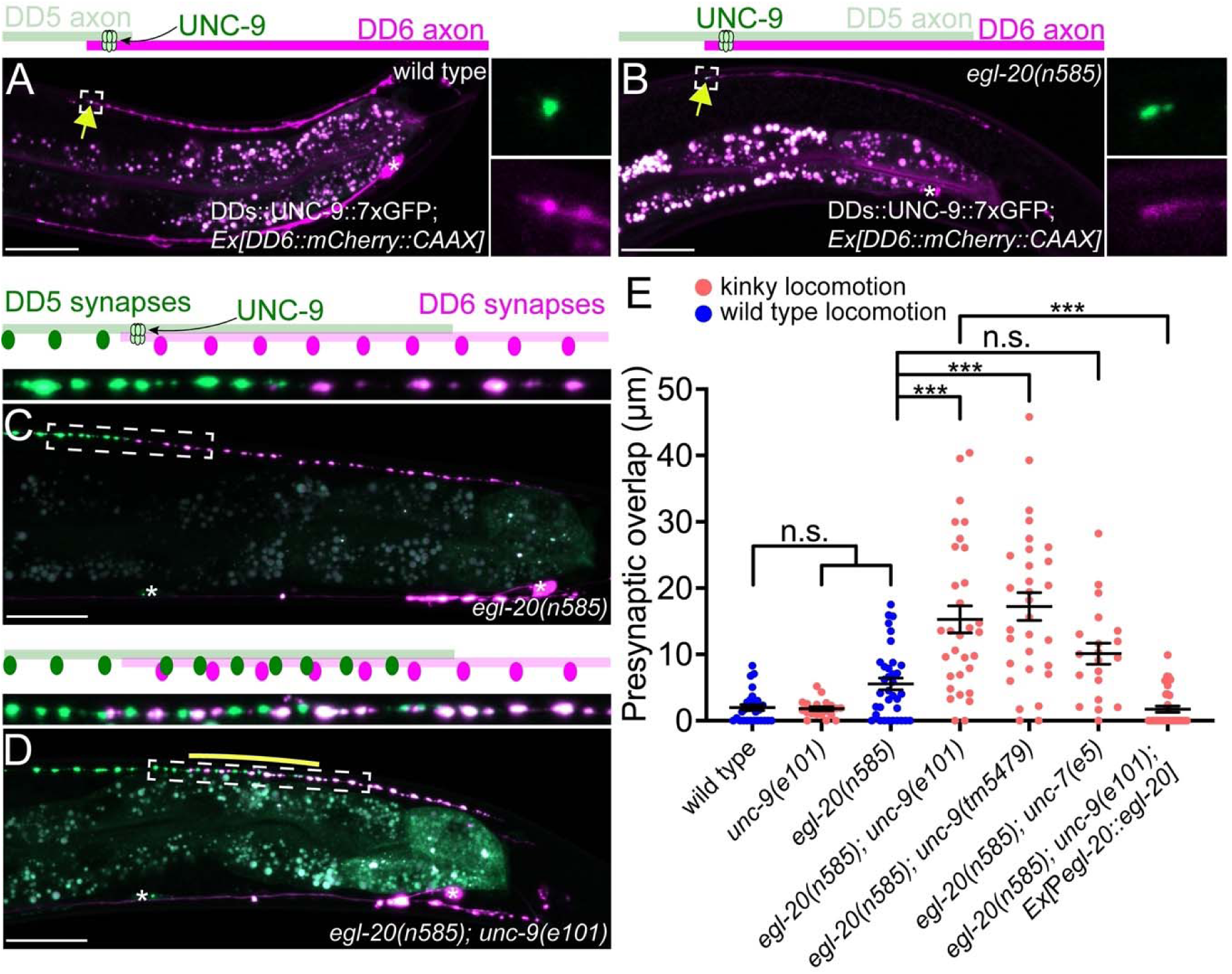
UNC-9/INX is localized at the presynaptic tiling border and is required for presynaptic tiling between DD5 and DD6 neurons. (A-B) Representative image of UNC-9::7×GFP localization at the tip of the anterior DD6 axon (indicated by yellow arrows) in wild type (A) and *egl-20(n585)* mutants (B). The magnified UNC-9::7×GFP and mCherry::CAAX signals, represented by dotted box, are shown to the right of merged images. (C-D) Representative image of presynaptic tiling in the *egl-20(n585)* (C), and *egl-20(n585); unc-9(e101)* (D) mutants. The magnified straightened image of the presynaptic tiling border, represented by dotted box, is shown above. Asterisks: DD5 and DD6 cell bodies. Scale bar: 20µm. (E) Quantification of presynaptic overlap between DD5 and DD6. Each dot represents a single animal. Black bars indicate mean ± SEM. n.s.: not significant; ***p<0.001.

We next attempted to understand the mechanisms of UNC-9 localization in the DD neurons by examining the mutants of known regulators of gap junction localization. In the nerve ring, clustered UNC-9 localization depends on NLR-1/CASPR (Meng and Yan, 2020). In vertebrates, ZO-1 tight junction protein plays crucial roles in gap junction plaque formation. For example, a recent work in zebrafish showed that loss of ZO1b resulted in a loss of Cx35.5 and Cx34.1 localization at the club ending synapses (Lasseigne et al., 2021). In the *nlr-1(gk366849: Q280Stop)* and *zoo-1(tm4133)* null mutants, we found that UNC-9::7×GFP localization appeared to be unaffected (**Supplemental figure 2-1**). We also observed normal UNC-9::7×GFP localization in the mutants of neuronal kinesin, *unc-104(e1265)*, in which axonal transport is severely compromised (Hall and Hedgecock, 1991) (**Supplemental figure 2-1**). It is therefore likely that the localization of UNC-9 at the axonal tiling border is controlled by previously unknown mechanisms.

### *unc-9* is required for presynaptic tiling between DD5 and DD6

Given that the presynaptic tiling pattern and UNC-9 gap junction localization are unaltered in *egl-20(n585)* mutants, we hypothesized that UNC-9 localized at the DD5 and DD6 presynaptic tiling border regulates presynaptic tiling. Consistent with this hypothesis, we observed presynaptic tiling defect in the double mutants of *egl-20(n585)* and *unc-9(e101* or *tm5479)* null mutants. Ectopic DD5 presynaptic puncta are formed in the DD5 posterior axon that overextended into the DD6 axonal region, which creates intermingled patterning of DD5 and DD6 presynaptic puncta (**Figures 2C-E**). The presynaptic tiling defect in the *egl-20(n585); unc-9(e101)* mutants is fully rescued by the P*egl-20::egl-20* transgene, which rescues the axonal tiling defect **(Supplemental figure 1-2)**, suggesting that overextension of DD5 posterior axon is necessary for observing the effect of *unc-9* in presynaptic patterning (**Figure 2E**). Consistently, *unc-9(e101)* single mutants did not exhibit presynaptic tiling defect due to the normal axonal tiling in the presence of *egl-20* (**Supplemental figure 1-2**). Importantly, loss of *unc-9* does not enhance the axonal tiling defect of *egl-20(n585)* mutants (**Supplemental figure 1-2**). Therefore, the presynaptic tiling defect in the *egl-20(n585); unc-9(e101)* double mutants is not due to an increased axonal tiling defect.

In order to determine whether the ectopic presynaptic RAB-3 puncta in *egl-20(n585); unc-9(e101)* mutants represent bona fide synapses, we examined co-localization between mCherry::RAB-3 expressed under the *flp-13* promoter and endogenously tagged NLG-1::GFP/Neuroligin, which is localized at the postsynaptic sites of the GABAergic neuromuscular junctions (Maro et al., 2015; McDiarmid et al., 2020; Tu et al., 2015). The ectopically formed mCherry::RAB-3 puncta at the posterior DD5 axon of *egl-20(n585); unc-9(e101)* mutants are apposed by NLG-1::GFP (**Supplemental figure 2-2**). This suggests that the ectopic RAB-3 puncta from DD5 neuron in *egl-20(n585); unc-9(e101)* double mutant animals likely represent bona fide synapses.

We also tested whether *unc-7*, another INX expressed in the DD neurons, also plays a role in presynaptic patterning. However, the degree of presynaptic tiling defect of *egl-20(n585); unc-7(e5)* double mutants is not significantly different from that of *egl-20(n585)* single mutants, although there is a tendency for a slightly larger overlap (**Figure 2E**). We therefore conclude that *unc-9* is the major INX that controls presynaptic tiling of DD5 and DD6.

DD neurons undergo remodeling at the end of the first larval stage, when the dorsal neurites switch their fate from dendrite to axon (White et al., 1978). We next asked whether EGL-20 and UNC-9 are required for establishing or maintaining axonal and presynaptic tiling, respectively. To test this, we examined axonal and presynaptic tiling at the second larval L2 stage when the DD remodeling is completed. Both axonal and presynaptic tiling were defective at the L2 stage in *egl-20(n585)* and *egl-20(n585); unc-9(e101)*, respectively (**Supplemental figure 2-3**). This suggests that *egl-20* and *unc-9* are required for the establishment of axonal and presynaptic tiling, respectively.

### UNC-9 functions cell autonomously in the DD neurons to control presynaptic tiling

*unc-9* is expressed broadly in the nervous system and non-neuronal tissues, including the DD neurons and their postsynaptic body wall muscles (Yeh et al., 2009). We therefore asked in which cells UNC-9 functions to control presynaptic tiling between DD5 and DD6 by conducting tissue-specific rescue experiments. Pan-neuronal expression of *unc-9* under the *rgef-1* promoter rescued the presynaptic tiling defect of the *egl-20(n585); unc-9(e101)* double mutants (**Figures 3A, C)**. Similarly, *unc-9* expression in the presynaptic DD neurons using the *flp-13* promoter but not in the postsynaptic body wall muscles using the *myo-3* promoter rescued the presynaptic tiling defect (**Figure 3B and 3D**). These results suggest that *unc-9* controls tiled presynaptic patterning in the presynaptic DD neurons. The slightly weaker rescue activity of P*flp-13::unc-9* compared with P*rgef-1::unc-9* may be due to a weaker expression of *unc-9* in the DD6 neuron. Expression of *unc-9* specifically in DD6 did not rescue the presynaptic tiling defect between DD5 and DD6 (**Figure 3C**). These results suggest that *unc-9* is required in both DD5 and DD6 neurons to control tiled presynaptic patterning between DD5 and DD6.

**Figure 3:**
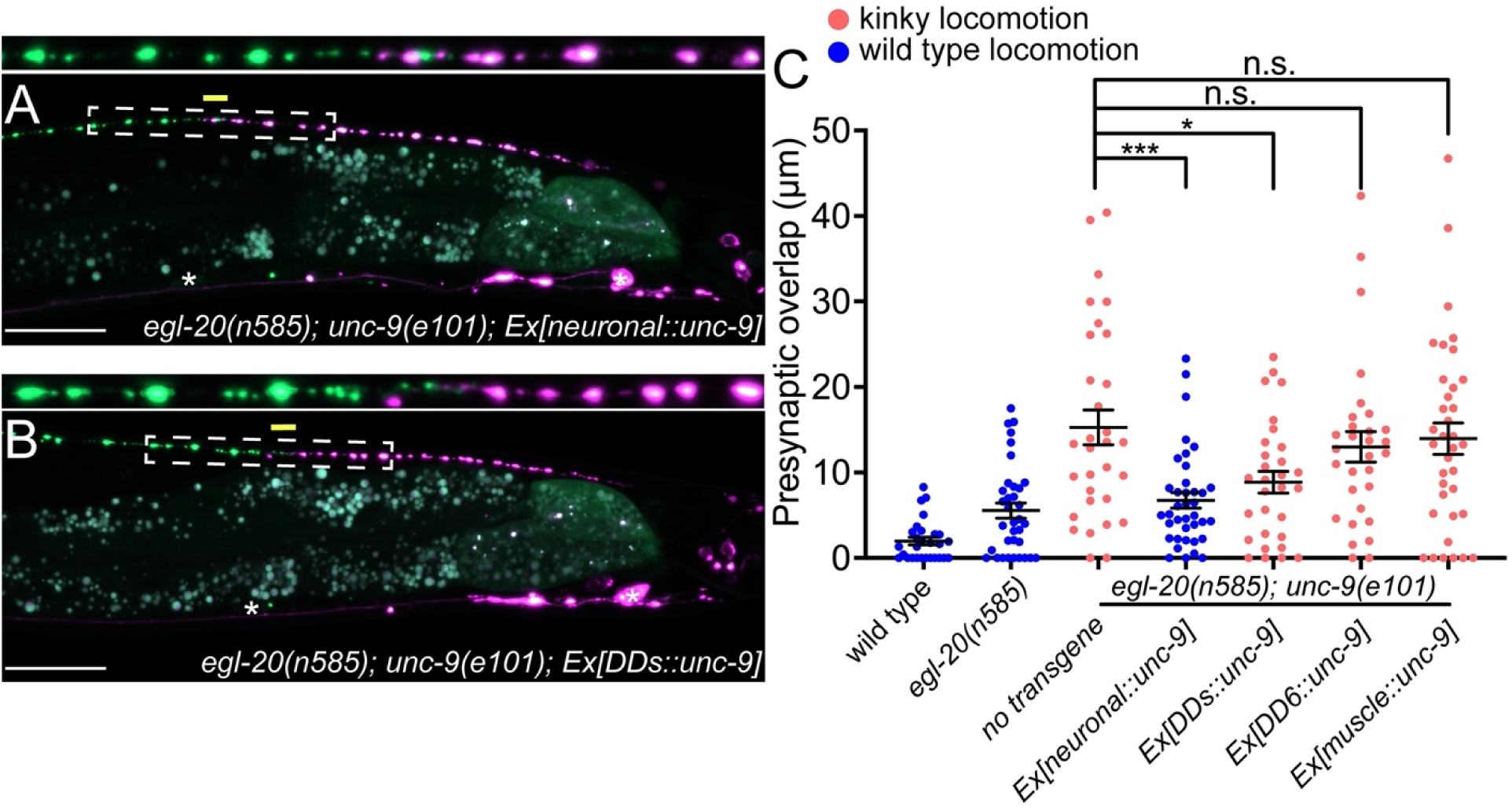
UNC-9/INX functions cell-autonomously to control presynaptic tiling between DD5 and DD6 neurons. (A-B) Representative image of presynaptic tiling in the *egl-20(n585); unc-9(e101)* double mutants with pan-neuronal expression of *unc-9* (A), and DD-specific expression of *unc-9* (B). The magnified straightened image of the presynaptic tiling border, represented by dotted box, is shown above. Asterisks: DD5 and DD6 cell bodies. Scale bar: 20µm. (C) Quantification of presynaptic overlap between DD5 and DD6. Each dot represents a single animal. Black bars indicate mean ± SEM. n.s.: not significant; *p<0.05; ***p<0.001.

### *unc-1/stomatin* is not required for the presynaptic tiling between DD5 and DD6

As UNC-9 forms gap junctions between DD5 and DD6, we next tested if UNC-9 controls tiled presynaptic patterning via its gap junction channel activity. It has been shown that UNC-1/stomatin is essential for opening the UNC-9 gap junction channels (Chen et al., 2007; Jang et al., 2017). In the loss-of-function mutants of *unc-1,* UNC-9 gap junction channel activity is completely abolished (Chen et al., 2007). Due to the defective UNC-9 gap junction channel activity, *unc-1* mutants exhibit a kinky locomotion (kinker) phenotype similar to *unc-9* mutants (**Supplemental Figure 3**). To test whether UNC-9’s function in regulating presynaptic tiling depends on its gap junction channel activity, we examined the presynaptic patterning of DD5 and DD6 in the *egl-20(n585); unc-1(e719)* double mutants. Interestingly, unlike *egl-20(n585); unc-9(e101)* double mutants, *egl-20(n585); unc-1(e719)* double mutants did not exhibit presynaptic tiling defect (**Figure 4A and 4D**). While UNC-9 gap junction activity is completely abolished in *unc-1* mutants, it is possible that another stomatin can mediate the UNC-9 gap junction activity in the DD neurons. We therefore examined *unc-24/stomatin,* as *unc-24* mutants also exhibit the kinker phenotype similar to *unc-1* and *unc-9* mutants (**Supplemental figure 3)**. Similar to *egl-20(n585); unc-1(e719)* mutants, *egl-20(n585) unc-24(miz225)* double mutants and *egl-20(n585) unc-24(miz226); unc-1(e719)* triple mutants did not exhibit presynaptic tiling defect (**Figure 4B, 4C and 4D**). These results raised the possibility that *unc-9*’s function in controlling presynaptic tiling between DD5 and DD6 is independent of its gap junction channel activity.

**Figure 4:**
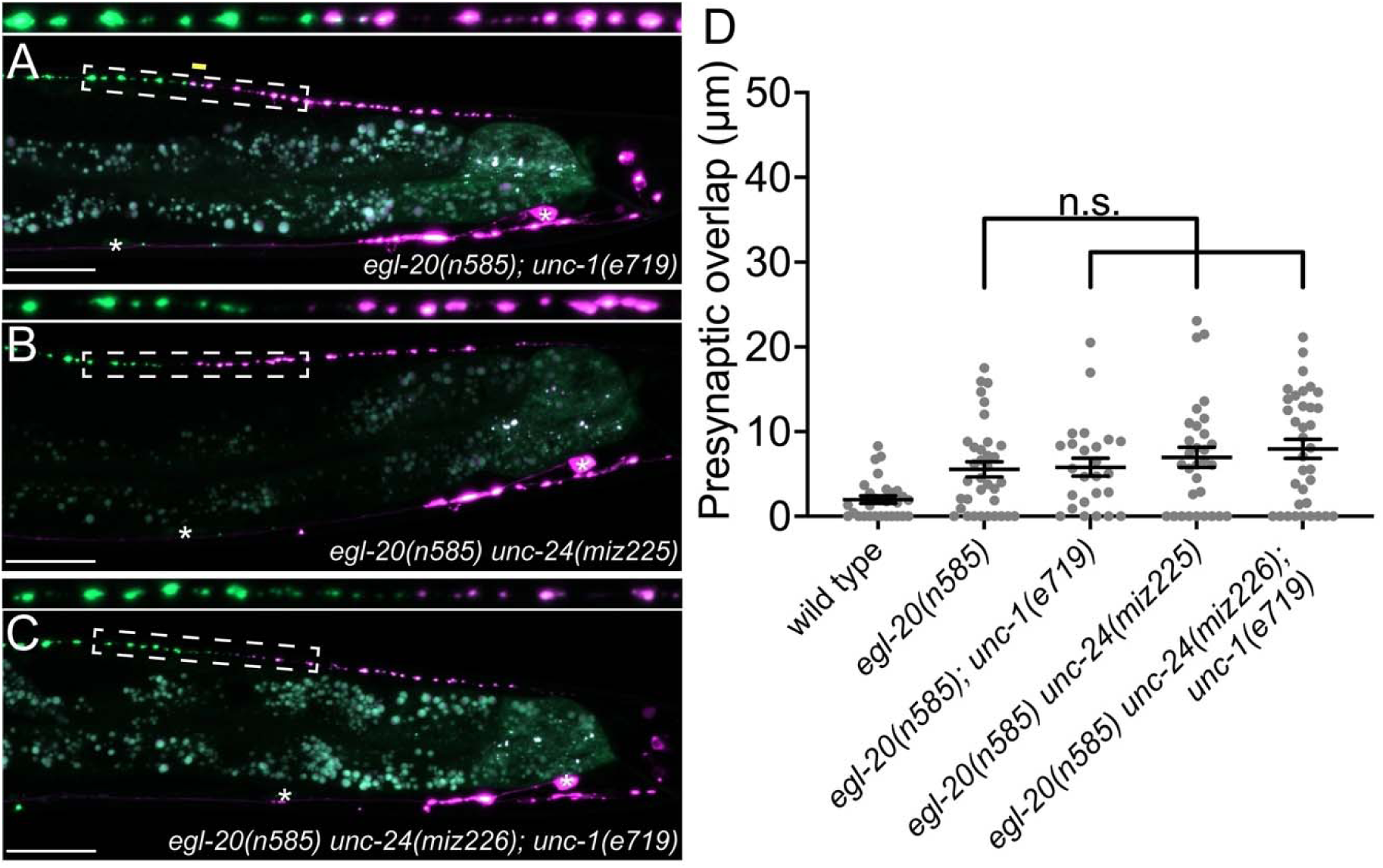
*unc-1/stomatin* is not required for presynaptic tiling between DD5 and DD6 neurons. (A-B) Representative image of presynaptic tiling in the *egl-20(n585); unc-1(e719)* double mutants (A), *egl-20(n585) unc-24(miz225)* (B), and *egl-20(n585) unc-24(miz226); unc-1(e719)* (C). The magnified straightened image of the presynaptic tiling border, represented by dotted box, is shown above. Asterisks: DD5 and DD6 cell bodies. Scale bar: 20µm. (D) Quantification of presynaptic overlap between DD5 and DD6. Each dot represents a single animal. Black bars indicate mean ± SEM. n.s.: not significant.

### Putative constitutively closed form of UNC-9 can control presynaptic tiling

While Cxs and INXs function primarily as gap junction channels, recent studies have revealed channel-independent roles for gap junction proteins (Dbouk et al., 2009; Elias et al., 2007; Kameritsch et al., 2013; Kameritsch et al., 2012; Miao et al., 2020; Olk et al., 2009). For example, *Drosophila* INXs (Inx2/3/4) control border cell migration independent of their gap junction channel activity (Miao et al., 2020). Cx43 and Cx26 mediate the migration of radial glia cells in a channel-independent manner (Elias et al., 2007). Given that *unc-1* is dispensable for *unc-9’s* function in presynaptic tiling, we tested if UNC-9 controls presynaptic tiling independent of its gap junction channel activity by using mutant UNC-9 with defective channel activity. Previous cryo-EM studies using *C. elegans* INX-6/INX have shown that a mutant INX-6 carrying an 18 amino acids deletion in its amino-terminal intracellular domain forms a constitutively closed gap junction channels (Burendei et al., 2020; Oshima et al., 2016a; Oshima et al., 2016b). In an attempt to generate a constitutively closed UNC-9 gap junction channel, we first examined the subcellular localization of UNC-9(ΔN18), which lacks the first 18 amino acids in its intracellular domain. Similar to UNC-9::GFP, UNC-9(ΔN18)::GFP expressed in the DD neurons was localized at the tip of DD6 anterior axon, the putative gap junction site between DD5 and DD6 (**Figure 5A and 5B**). This result indicates that the 18 amino acid deletion does not alter either the synthesis or subcellular localization of the UNC-9 protein. We then tested if the putative channel-defective *unc-9(ΔN18)* can rescue the presynaptic tiling defect by expressing it under the *rgef-1* pan-neuronal promoter or under the *flp-13* DD-neuron-specific promoter. Surprisingly, both pan-neuronal and DD neuron-specific expression of *unc-9(Δ18N)* rescued the presynaptic tiling defect of *egl-20(n585); unc-9(e101)* mutants (**Figure 5D**). Importantly, pan-neuronal expression of wild type *unc-9* but not *unc-9(ΔN18)* rescued the kinker phenotype of the *unc-9* mutant. As the locomotion defects of *unc-9* mutants is largely due to the defective gap junction channel activity (Kawano et al., 2011), the failure to rescue the locomotion defects of *unc-9* mutants by *unc-9(Δ18N)* strongly suggests that UNC-9(ΔN18) forms a defective, likely a constitutively closed, gap junction channel.

**Figure 5:**
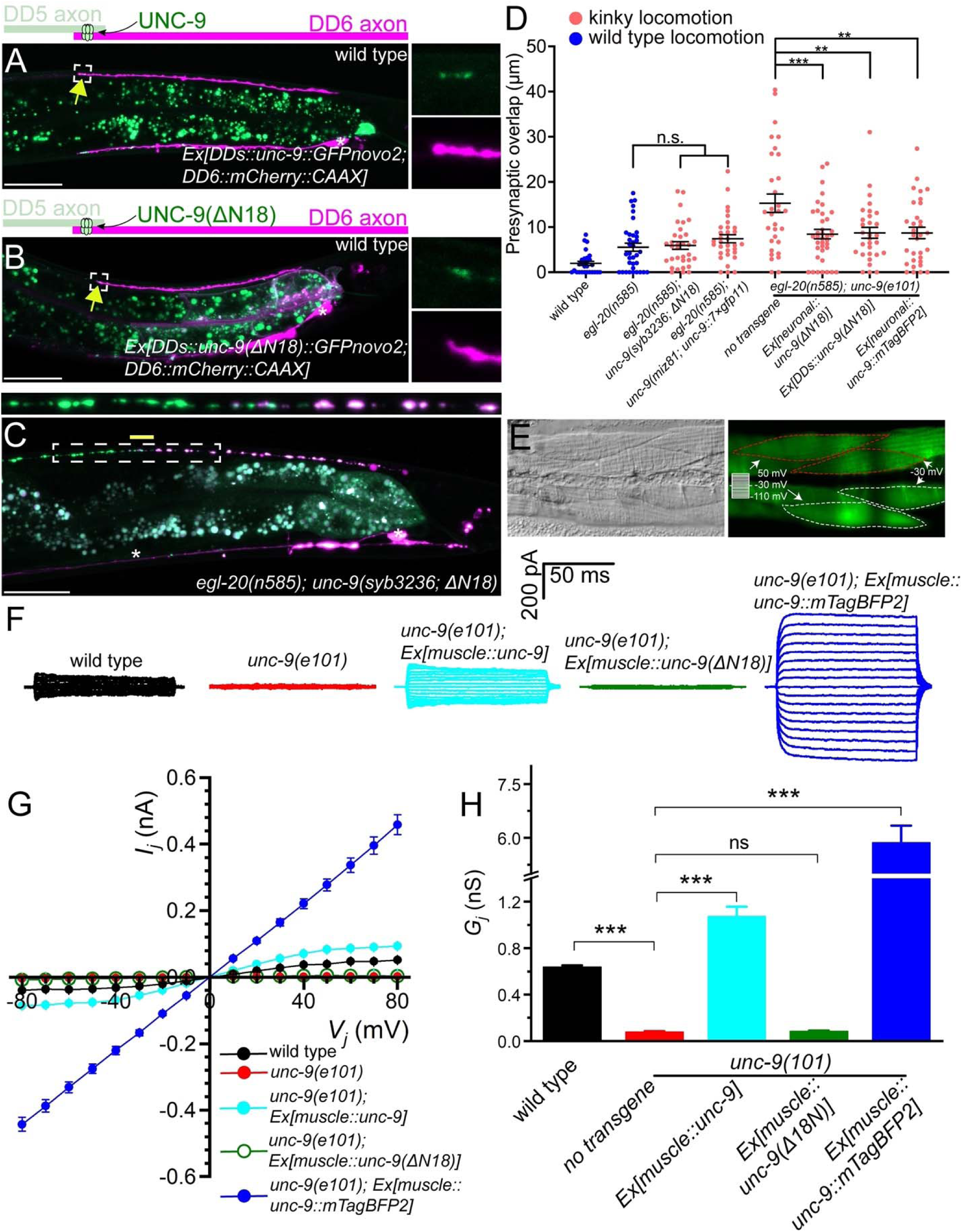
UNC-9 gap junction channel activity is dispensable for its function in presynaptic patterning. (A-B) Representative image of UNC-9::GFP (A) and UNC-9(ΔN18)::GFP (B) localization at the tip of the anterior DD6 axon (indicated by yellow arrows) in wild type. The magnified UNC-9::7×GFP and mCherry::CAAX signals, represented by dotted box, are shown to the right of merged images. (C) Representative image of presynaptic tiling in the *egl-20(n585); unc-9(syb3236; ΔN18)* double mutants. The magnified straightened image of the presynaptic tiling border, represented by dotted box, is shown above. Asterisks: DD5 and DD6 cell bodies. Scale bar: 20µm. (D) Quantification of presynaptic overlap between DD5 and DD6. Each dot represents a single animal. Black bars indicate mean ± SEM. n.s.: not significant; **p<0.01; ***p<0.001. (E) Image of adjacent body wall muscles labelled with GFPnovo2. (F) Intra-quadrant coupling between a pair of neighboring L1-L2 or R1-R2 cells in different mutants. (G) Quantification of the junctional currents (*I_j_*). (H) Quantification of the junctional conductance (*G_j_*).

To directly assess the effect of the ΔN18 mutation on UNC-9 gap junction channel activity, we tested whether expression of UNC-9(ΔN18) can rescue the electrical coupling defect of *unc-9(e101)* mutant. In these experiments, we expressed either wild type *unc-9* or *unc-9(ΔN18)* in the muscle cells using the muscle-specific *myo-3* promoter (P*myo-3*) in the *unc-9(e101)* mutant background. Cytoplasmic GFP was co-expressed in the muscle cells of these worms to serve as a transformation marker. We then performed dual whole-cell voltage clamp recordings on two contiguous neighboring ventral muscle cells from two different rows of muscle cells within the same quadrant (**Figure 5E**). Junctional currents (*I_j_*) were measured from one muscle cell held constant at −30 mV while a series of membrane voltage steps (−110 to +50 mV at 10-mV intervals) were applied to the other neighboring muscle cell of the pair from a holding voltage of −30 mV (**Figure 5E**). We plotted the *I_j_* and transjunctional voltage (*V_j_*) relationships and quantified the junctional conductance (*G_j_*) from the slope of the linear portion of the *I_j_* -*V_j_*curves. While *I_j_* was prominent in the wild type, *unc-9(e101)* mutant showed very little *I_j_* (**Figure 5G**), similar to the *unc-9(fc16)* null mutants analyzed in our earlier studies (Liu et al., 2011; Liu et al., 2006). As expected, wild type UNC-9 restored the junctional current in the *unc-9(e101)* mutants, while *unc-9(e101)* mutants expressing *unc-9(ΔN18)* completely lacked the intra-quadrant coupling similar to *unc-9(e101)* mutants (**Figure 5F and 5H**). These results confirm that UNC-9(ΔN18) is unable to form functional gap junction channels, while maintaining the ability to control tiled presynaptic patterning between DD5 and DD6.

The successful rescue of the presynaptic tiling defect by the *unc-9(ΔN18)* transgenes could be due to the nature of overexpression of the transgene from extra-chromosomal concatemer arrays. To exclude this possibility, we generated *unc-9(ΔN18)* mutants using CRISPR/Cas9 genome editing. Similar to *unc-9(e101)* mutants*, unc-9(syb3236; ΔN18)* mutant animals exhibit kinker phenotype (**Supplemental Figure 3**), suggesting that UNC-9(ΔN18) does not form functional gap junction channels. Despite the kinker phenotype, *egl-20(n585); unc-9(ΔN18)* double mutants did not exhibit presynaptic tiling defect between DD5 and DD6 (**Figures 5B and 5C**). This further confirmed that UNC-9 does not require its gap junction channel activity to controls presynaptic tiling.

### Putative constitutively open form of UNC-9 can control presynaptic tiling

Previously, we showed that an UNC-9::GFP fusion protein may form constitutively open gap junction channels in muscles and that such gap junctions do not require UNC-1 to function (Chen et al., 2007). The dispensability of UNC-1 to UNC-9::GFP gap junctions has also been shown with *C. elegans* neurons (Chen et al., 2007; Jang et al., 2017). To further substantiate the potential channel-independent function of UNC-9 in controlling presynaptic tiling, we examined whether *unc-9::mTagBFP2* can rescue the presynaptic tiling defect. We chose UNC-9::mTagBFP2 instead of UNC-9::GFP, as the GFP signal interferes with our presynaptic marker (GFP::RAB-3). We first tested whether UNC-9::mTagBFP2 forms an open gap junction channel similar to UNC-9::GFP by testing whether it can rescue the electrical coupling defects of *unc-9(e101)* mutants. The muscle expression of *unc-9::mTagBFP2* in *unc-9(e101)* mutant resulted in a much larger *I_j_* and *G_j_* than those obtained with wild type *unc-9* (**Figure 5F-H**), which is reminiscent of our previous observation with *unc-9::GFP* (Chen et al., 2007). We then expressed *unc-9::mTagBFP2* pan-neuronally using the *rgef-1* promoter, and tested whether it has the ability to control presynaptic tiling patterning. Similar to *unc-9(ΔN18)*, pan-neuronal expression of *unc-9::mTagBFP2* rescued the presynaptic patterning defect of *egl-20(n585); unc-9(e101)* mutants (**Figure 5C**). Importantly, pan-neuronal expression of *unc-9::mTagBFP2* did not rescue the kinker phenotype of *egl-20(n585);unc-9(e101)* mutants (**Supplemental Figure 3**), which is probably because hyperactive gap junction channel activity formed by UNC-9::mTagBFP2 impaired rather than restored the function of the locomotor neural circuit. We further confirmed this result by examining the presynaptic tiling pattern in *egl-20(n585); unc-9(miz81; unc-9::7×gfp11)* mutants. UNC-9::7×GFP11, which by itself is not fluorescent, has seven tandem repeats of GFP11 at the carboxy-terminus of UNC-9 and hence is expected to have a similar effect on gap junction function as UNC-9::GFP or UNC-9::mTagBFP2. While UNC-9::7×GFP localizes to the putative gap junction sites between DD5 and DD6 axons, *unc-9(miz81; unc-9::7×gfp11)* mutants exhibit a kinker phenotype (**Supplemental Figure 3**), suggesting that they form defective, likely an open form of gap junctions. Despite the kinker phenotype, *egl-20(n585); unc-9(miz81; unc-9::7×gfp11)* mutants do not exhibit the presynaptic tiling defect (**Figure 5C**). These results suggest that an increased activity of UNC-9 gap junctions does not compromise the physiological role of UNC-9 in controlling the presynaptic tiling pattern. Taken together, our results indicate that UNC-9 controls presynaptic tiling pattern through a gap junction channel-independent function.

## Discussion

Here we have established a novel system to study neuronal and synaptic tiling in the DD type GABAergic motor neurons in *C. elegans*. We found that EGL-20/Wnt negatively regulates the length of the posterior axon and dendrite of DD5. We showed that presynaptic tiling requires UNC-9/INX, while the position of postsynaptic spines appears to depend on the length of the posterior dendrite. UNC-9 is localized at the presynaptic tiling border between DD5 and DD6 where it forms gap junction channels. Strikingly, the gap junction channel activity of UNC-9 is dispensable for its function in establishing presynaptic tiling.

### Redundant actions of Wnt and INX in establishing DD presynaptic tiling

In wild type animals, the presynaptic domain of each DD neuron does not overlap with those from neighboring DDs, because of their tiled axonal patterning. However, our observation revealed that the axonal tiling is dispensable for the presynaptic tiling, which is controlled by UNC-9. Therefore, the presynaptic tiling of the DD neurons is governed by two redundant pathways: Wnt-dependent axonal tiling and INX-dependent presynaptic tiling. This observation is very unique compared to other systems such as the L1 lamina neuron in *Drosophila* where axonal tiling defects cause disrupted synaptic connections (Millard et al., 2007). DD neurons are critical for the coordinated contractions and relaxations between the dorsal and ventral body wall muscles during sinusoidal locomotion of the worms (Kawano et al., 2011). As the synapse is the functional unit for this functionality of the DD neurons, the redundant mechanisms to tile synapses in DD neurons by Wnt and INX may ensure the proper functions of the DD neurons. This may be because the DD neurons are the only GABAergic-class of motor neurons in the dorsal nerve cord and thus require more robust control of their synaptic patterning.

The degree of presynaptic tiling defect of *egl-20(n585); unc-9(e101)* mutants is smaller than the degree of axonal tiling defect, suggesting the presence of additional factors that regulate presynaptic patterning between DD5 and DD6 along with *unc-9.* Previously, we showed that spatial expression pattern of two Wnt proteins (LIN-44 and EGL-20) defines topographic presynaptic patterning of DA8 and DA9 in the absence of Plexin-dependent presynaptic tiling mechanism (Mizumoto and Shen, 2013b). It is therefore possible that LIN-44/Wnt contributes towards the positioning of DD5 synapses in the *egl-20; unc-9* mutants. We could not test this hypothesis as the expression pattern of the DD5/DD6 presynaptic tiling marker is disrupted in *lin-44* mutant.

### Mechanisms of INX-dependent synapse patterning and potential links to synaptopathies

How does UNC-9 control presynaptic tiling between DD5 and DD6? UNC-9 gap junctions are localized at the presynaptic tiling border in both wild type and in *egl-20* mutants. The specific subcellular localization of UNC-9 gap junctions at the presynaptic tiling border suggests that UNC-9 acts as a positional cue by forming a highly localized gap junction plaque that locally restricts synapse formation. Indeed, loss of *unc-9* results in the ectopic synapse formation in the distal posterior axonal region of DD5. Consistent with our observations, recent work in mouse cortical neurons showed that PANX1 inhibits synapse formation (Sanchez-Arias et al., 2020; Sanchez-Arias et al., 2019). It is therefore possible that gap junction proteins have conserved functions as negative regulators for synapse formation. Previously, UNC-9 was shown to localize at the nonjunctional perisynaptic region in the DD neurons to regulate active zone differentiation, possibly as hemichannels (Yeh et al., 2009). *unc-9*’s function in promoting presynaptic assembly contrasts with our findings that *unc-9* restricts synapse formation. It is possible that distinct localization of UNC-9 may inhibit or promote synapse formation through distinct downstream molecular pathways. While we do not know how UNC-9 locally restricts synapse formation, it is possible that it acts through actin cytoskeleton and its regulators, as proper actin cytoskeleton formation is essential for the synapse formation (Aiken and Holzbaur, 2021; Hendi et al., 2019). Cx proteins have been shown to interact with various actin cytoskeletal regulators including Rap1, IQGAP, ZO-1, Claudin, and Drebrin (Falk et al., 2014; Lasseigne et al., 2021; Olk et al., 2009). Indeed, we have previously shown that RAP-2 small GTPase and its effector kinase MIG-15/TNIK are crucial for the presynaptic tiling of the DA8 and DA9 neurons (Chen et al., 2018). However, we did not observe DD5/DD6 presynaptic tiling defects in the *rap-1/Rap1, rap-2/Rap2, pes-7/IQGAP, zoo-1/ZO-1, vab-9/Caludin, dbn-1/Drebrin* and *mig-15/TNIK* in the *egl-20* mutant background (data not shown). The DD5 and DD6 neurons seem to control presynaptic tiling through an uncharacterized mechanism.

Mutations in gap junction proteins are often associated with various synaptopathies caused by abnormal synapse number and position (Lapato and Tiwari-Woodruff, 2018). The present work that revealed novel function of a gap junction protein in presynaptic patterning will help us better understand how mutations in gap junction proteins cause synaptopathies. Further candidate and forward screenings are essential to uncover the molecular mechanisms that underlie INX-dependent synapse patterning.

### Limitation of the present work

This work showed that the presynaptic tiling of the DD neurons is controlled by Wnt-dependent axonal tiling and UNC-9-dependent presynaptic tiling. The redundant mechanisms to set up tiled presynaptic arrangement argues that it is important for the function of these neurons in locomotion. However, we could not test the effect of disrupted presynaptic tiling of the DD GABAergic motor neurons on locomotion, as all mutants with disrupted presynaptic tiling exhibited a kinker phenotype due to defective UNC-9-channel activity, which prevented us from examining the direct effect of presynaptic tiling defect on locomotion. Creating specific *unc-9* mutants which specifically disrupt UNC-9’s function in presynaptic tiling without affecting its channel activity will help us uncovering the functional importance of the presynaptic tiling of the DD neurons.

DD neurons undergo synaptic remodelling at the end of the first larval stage L1, when the dorsal dendrites switch their fates to axons to form presynaptic connections with the dorsal body wall muscles (White et al., 1978). We showed that at mid L2 stage, *egl-20(n585); unc-9(e101)* double mutants exhibit presynaptic tiling defects (**Supplemental Figure 2-2**). This suggests that *elg-20* and *unc-9* are required for the establishment of the axonal and presynaptic tiling, respectively. However, due to the small size of the L1 animals, we could not observe the presynaptic tiling patterning during DD remodeling to observe how the presynaptic tiling is established. Further work is required to determine whether *unc-9* is also required throughout the lifespan of animals to maintain presynaptic tiling.

We did not identify UNC-9’s downstream effectors (see above), nor the upstream components that are required for the UNC-9 localization at the presynaptic tiling border. There are several genes that are necessary for the proper clustering of gap junction channels. However, UNC-9 localization was unaffected in the mutants of *zoo-1/ZO-1* and *nlr-1/CASPR*. Recent work showed that cAMP-dependent axonal transport is required for the proper localization of UNC-9 in the VA-class of cholinergic motor neuron in *C. elegans* (Palumbos et al., 2021). While we observed normal UNC-9 in the *unc-104/Kif1A* mutants, it is possible that UNC-9 transport is regulated by another motor protein in the DD neurons. The stereotypical localization of UNC-9 at the presynaptic border between DD neurons provides an ideal platform to carry out genetic screens to decipher the molecular mechanisms that underlie gap junction localization.

## Materials and methods

### Strains

Bristol N2 strain was used as wild type reference. All strains were cultured in the nematode growth medium (NGM) with OP50 as described previously (Brenner, 1974) at 25°C. The following alleles were used in this study: *egl-20(n585), unc-9(e101), unc-9(tm5479), unc-1(e719), unc-9(miz81; unc-9::7×gfp11), unc-9(syb3236; ΔN18), unc-24(miz225), unc-24(miz226), unc-104(e1265), nlr-1(gk366849), zoo-1(tm4133), nlg-1(yv15), unc-7(e5)*.

Genotyping primers are listed in the supplemental material.

### Transgenes

The transgenic lines with extrachromosomal arrays were generated using the standard microinjection method (Fire, 1986; Mello et al., 1991). The integration of the extrachromosomal arrays into the chromosomes was conducted by standard UV irradiation method.

The following transgenes were used in this study: *wyIs442 (*P*flp-13::2×GFP::rab-3,* P*plx-2::2×mCherry::rab-3); wyIs486 (*P*flp-13::2×GFP,* P*plx-2::2×mCherry); juIs463 (*P*flp-13::GFP1-10,* P*ttx-3::RFP); ufIs126 (*P*flp-13::acr-12::GFP); mizEx69, mizEx407 (*P*rgef-1::unc-9,* P*odr-1::GFP); mizEx72, mizEx420 (*P*rgef-1::unc-9(ΔN18),* P*odr-1::GFP); mizEx416 (*P*flp-13::unc-9,* P*odr-1::GFP); mizEx429, mizEx433 (*P*flp-13::unc-9(ΔN18),* P*odr-1::GFP); mizEx430 (*P*flp-13::mCherry::rab-3,* P*odr-1::GFP); mizEx510 (*P*myo-3::unc-9,* P*odr-1::GFP), mizEx496, mizEx497 (*P*myo-3::unc-9,* P*myo-3::GFPnovo2,* P*odr-1::GFP); mizEx499, mizEx500 (*P*myo-3::unc-9(ΔN18),* P*myo-3::GFPnovo2,* P*odr-1::GFP); mizEx498, mizEx499 (*P*myo-3::unc-9(ΔN18),* P*myo-3::GFPnovo2,* P*odr-1::GFP); mizEx500, mizEx501 (*P*myo-3::unc-9::mTagBFP2,* P*myo-3::GFPnovo2,* P*odr-1::GFP); mizEx515 (*P*egl-20::egl-20,* P*odr-1::GFP)*.

### Key Strains

UJ1044 *egl-20(n585); wyIs442*

UJ1215 *egl-20(n585); unc-9(e101); wyIs442*

UJ1543 *egl-20(n585); unc-9(syb3236; ΔN18); wyIs442*

UJ1261 *unc-9(miz81; unc-9::7×gfp11); juIs463*

### Plasmid construction

*C. elegans* expression clones were made in a derivative of pPD49.26 (A. Fire), the pSM vector (a kind gift from S. McCarroll and C. I. Bargmann). *unc-9* cDNAs were amplified with Phusion DNA polymerase (NEB) from N2 cDNA library synthesized with Superscript III first-strand synthesis system (Thermo Fisher Scientific) The amplified cDNAs were cloned into the AscI and KpnI sites of pSM vector using Gibson assembly method (Gibson, 2011)

### Bashed *plx-2* promoter construction for DD6-specific expression

A 4.5 Kb PCR fragment of the 5’ promoter region of *plx-2* was cloned upstream of GFP between *Hinc*II and *Msc*I in pPD95.75, to generate pRI20. A sub-fragment of the *plx-2* promoter that drives selective expression in the DD6 neuron was generated by deleting a 3587 bp internal region in the plx-2 promoter of pRI20, by digesting with *Hpa*I and *Bam*HI, end-filling the *Bam*HI 5’ overhang with Klenow and blunt-end ligation to generate pRI50. The resulting *plx-2* promoter was re-cloned into the *Sph*I and *Asc*I sites of the pSM vector.

### CRISPR

The *loxP::myo-2::NeoR* dual-selection cassette vector (Au et al., 2019; Norris et al., 2015) was used to construct the repair template plasmids for *unc-9(miz81; unc-9::7×gfp11)*. *7×gfp11* sequence was cloned into the *Sac*II site of the *loxP::myo-2::NeoR* plasmid using Gibson assembly (Gibson, 2011) The 5’ and 3’ homology arms of *unc-9* repair template were amplified from N2 genomic DNA using Phusion DNA polymerase and cloned into the *Sac*II and *Not*I restriction sites, respectively, by Gibson assembly.

pTK73 plasmid was used as a backbone vector for gRNA expression plasmid construction for *unc-9(miz81; unc-9::7×gfp11)* strain (Obinata et al., 2018). The following guide RNA was used *unc-9#1:* AATTAAACCCCATTTCAGGA.

To generate *unc-9(miz81; unc-9::7×gfp11), unc-9* repair template plasmid, one *unc-9* sgRNA plasmid and Cas9 plasmid (Friedland et al., 2013) were co-injected into young adults. The screening of the genome edited animals was conducted as previously described (Au et al., 2019; Norris et al., 2015). F1 progenies were treated with Geneticin (G418) (Sigma-Aldrich) (50 mg/mL) for NeoR selection. Animals with uniform pharyngeal expression of P*myo-2::GFP* were selected as genome-edited candidates. Selection cassette was excised out by Cre recombinase (pDD104, Addgene #47551). Progenies without pharyngeal P*myo-2::GFP* expression were isolated. Successful genome edited candidate animals were confirmed via PCR and Sanger sequencing.

In order to generate *unc-24(miz225)* & *(miz226), unc-24* oligonucleotide primer repair template, and *unc-24* sgRNA and Cas9 ribonucleoprotein complex (Dokshin et al., 2018), were co-injected into young adults. Both *unc-24(miz225)* & *(miz226)* alleles share identical mutations.

The repair template included a mutation that replaced the PAM sequence to a thymine (T) base and an *Eco*RI restriction enzyme to create a premature stop codon and frameshift. Animals exhibiting the kinker phenotype were selected as genome edited candidates, and *Eco*RI site was used for genotyping the mutant alleles. Successful genome edited candidate animals were confirmed by PCR and Sanger sequencing.

*unc-24* gRNA: CGTTGAGCAGCGTTGCGAAA

Repair template oligos:

Forward:GGAAGGAGAGAACATGGGGATGTCTGCGTTGAGCAGCGTTGCGAAAgaattc**T**GATGCTGGTCAACAGTTGTGGCAAGTTATTGGACCAGTATTCG

Reverse:CGAATACTGGTCCAATAACTTGCCACAACTGTTGACCAGCATC**A**gaattcTTTC GCAACGCTGCTCAACGCAGACATCCCCATGTTCTCTCCTTCC

### Confocal and stereo microscopy

Images of fluorescently tagged fusion proteins used here (GFP and mCherry) were captured in live *C. elegans* using a Zeiss LSM800 Airyscan confocal microscope (Carl Zeiss, Germany) with oil immersion lens 40× magnification (Carl Zeiss, Germany). Worms were immobilized on 5% agarose pad using a 3:1 mixture of 0.225 M BDM (2,3-butanedione monoxime) (Sigma-Aldrich) and 7.5 mM levamisole (Sigma-Aldrich). Images were analyzed using Zen software (Carl Zeiss). Images were straightened with ImageJ (NIH, USA). eighteen to 24 Z-stack images were taken for each animal to encompass the cell bodies, axons, and synapses of the DD5 and DD6 neurons. L4.4–L4.6 larval stage animals, judged by the stereotyped shape of the developing vulva (Mok et al., 2015) were used for quantification. Stereoscope images were taken on Zeiss Stemi 305 with Zeiss Labscope.

### Statistics

Generated data were analysed and processed by Prism9 (GraphPad Software, USA). We applied used the one-way ANOVA method with post hoc Tukey’s multiple comparison test for comparison between three or more parallel groups with multiple plotting points. T-test was used for comparison between two binary points. Data were plotted with error bars representing standard errors of mean (SEM). *, **, and *** represent p-value <0.05, <0.01, <0.001, respectively.

### Primers

*egl-20(n585)* – wild type PCR product will be digested by *HpyCH*4V

Forward: 5’ CTCTTAAAAACTTACCTCTCAAATTTGAACTTATTCTTGC 3’

Reverse: 5’ CCTCATTACCATTCAACTGATAG 3’

*unc-24(miz225)* & *(miz226)* – mutant PCR product will be digested by *Eco*RI

Forward: 5’ CCACAGATCGTGGTCTCGTGGAAC 3’

Reverse: 5’ CTGACATTCGCTCCACCAAGTGTTTTAGC 3’

*nlr-1(gk366849)* – wild type PCR product will be digested by *Mfe*I

Forward: 5’ GTTTGCTCTCTTCATCAATCACTACATCC 3’

Reverse: 5’ CGCCATAAAACGATATATTATGTGTAG 3’

*zoo-1(tm4133)*

Mutant forward: 5’ CAGGTCGGCGGAAGTGTCGGAGTACGTG 3’

Wild type forward: 5’ CCGAATCAAGCGACCGCCGAGCAAATTGC 3’

Reverse: 5’ GTGCCAGCTGAAGACGTTCAACAGACTCG 3’

Electrophysiology

Adult (day 1) hermaphrodite animals were immobilized and dissected as described previously (Liu et al., 2011; Liu et al., 2006). Briefly, an animal was immobilized on a glass coverslip by applying Vetbond^TM^ Tissue Adhesive (3M Company, MN, USA). Application of the adhesive was generally restricted to the dorsal middle portion of the animal, allowing the head and tail to sway during the experiment. A longitudinal incision was made in the dorsolateral region. After clearing the viscera, the cuticle flap was folded back and glued to the coverslip, exposing the ventral nerve cord and the two adjacent muscle quadrants. A Nikon FN-1 microscope equipped with a 40× water-immersion objective and 15× eyepieces was used for viewing the preparation. Borosilicate glass pipettes with a tip resistance of 3∼5 MΩ were used as electrodes for voltage-clamp recordings in the classical whole-cell patch clamp configuration with a Multiclamp 700B amplifier (Molecular Devices, Sunnyvale, CA) and the Clampex software (version 11, Molecular Devices). To record *I_j_*, the membrane potential (*V_m_*) of both cells was held at −30 mV, from which a series of voltage steps (−110 mV to +50 mV at 10 mV intervals and 100 ms duration) were applied to one cell (Cell 1), whereas the other cell (Cell 2) was held constant to record *I_j_*. *V_j_*was defined as *V_m_* of Cell 2 minus *V_m_* of Cell 1. Series resistance was compensated to approximately 80% in the voltage-clamp experiments. Data was sampled at a rate of 10 kHz after filtering at 2 kHz.

## Acknowledgments

We would like to thank Cornelia I. Bargmann, Catharine H. Rankin and Michael M. Francis for sharing plasmids and strains, Shinsuke Niwa for sharing pTK73 gRNA plasmid, Atsunori Oshima for conducting preliminary experiments, Donald G. Moerman and the Mizumoto lab members for discussions. *unc-9(syb3236; ΔN18)* was generated by SunyBiotech, China. Some strains used in this study were obtained from the *Caenorhabditis* Genetics Center, CGC, funded by National Institute of Health (NIH) - Office of Research Infrastructure Programs (P40 OD010440), *C. elegans* gene knockout consortium, and the National Bioresource Project, Japan. This work is supported by CIHR AWD-017638 (K.M.), CIHR PJT-180563 (K.M.), NIH R01NS109388 (Z.W.), and NIH R01MH085927 (Z.W.). K.M. is a recipient of Canada Research Chair and Michael Smith Foundation for Health Research Scholar. A.H. is a recipient of NSERC CGS-D and the UBC 4-year fellowships.

---

**Supplemental figure 1-1:**
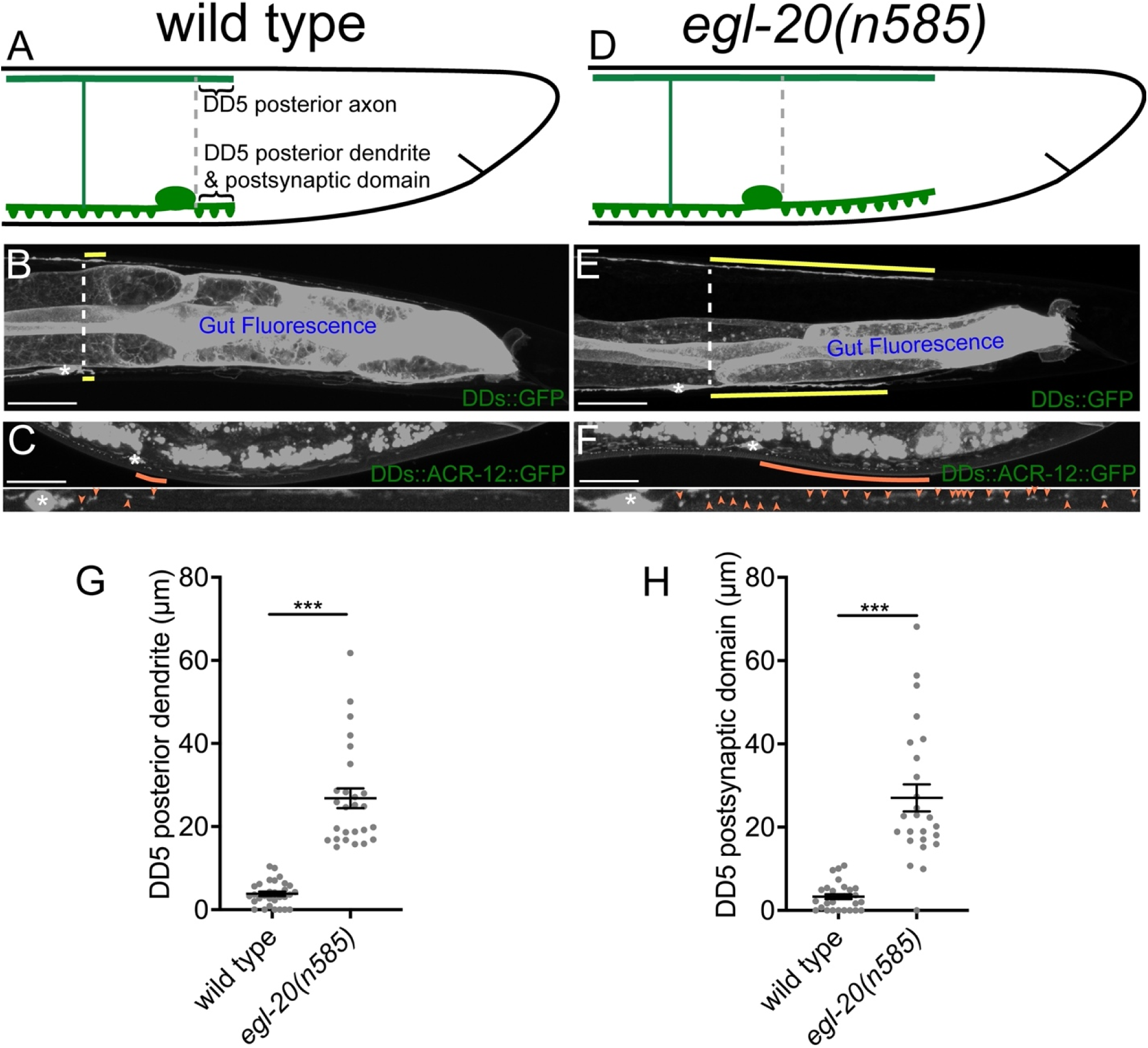
DD5 posterior dendrite and postsynaptic domain overextend posteriorly in the loss of *egl-20*. (A) Schematic of posterior axon and dendrite of DD5 neuron in wild type. Puncta represent postsynaptic sites. We defined the posterior axon of DD5 as follows: A vertical line was extended from the posterior end of the DD5 neuron cell body vertically to the dorsal nerve cord. The length of the posterior DD5 axon, dendrite and postsynaptic domains were measure from the posterior tip of the axons, dendrites and postsynaptic spine to the vertical line. (B) Representative image of posterior DD5 axon and dendrite in wild type in gray scale. (C) Representative image of ACR-12::GFP in the DD5 posterior dendrite of wild type in gray scale. (D) Schematic of posterior axon and dendrite of DD5 neuron in *egl-20(n585)* mutants. Puncta represent postsynaptic dendritic spines. (E) Representative image of posterior DD5 axon and dendrite in *egl-20(n585)* mutants in gray scale, posterior axon and dendrites represented by yellow line. (F) Representative image of ACR-12::GFP in the DD5 posterior dendrite of *egl-20(n585)* mutants in gray scale, posterior axon and dendrites represented by orange line. Asterisks: DD5 cell bodies. Scale bar: 20µm. (G) Quantification of DD5 posterior dendrite length. (H) Quantification of DD5 postsynaptic domain length. Each dot represents a single animal. Black bars indicate mean ± SEM. ***p<0.001.

**Supplemental figure 1-2:**
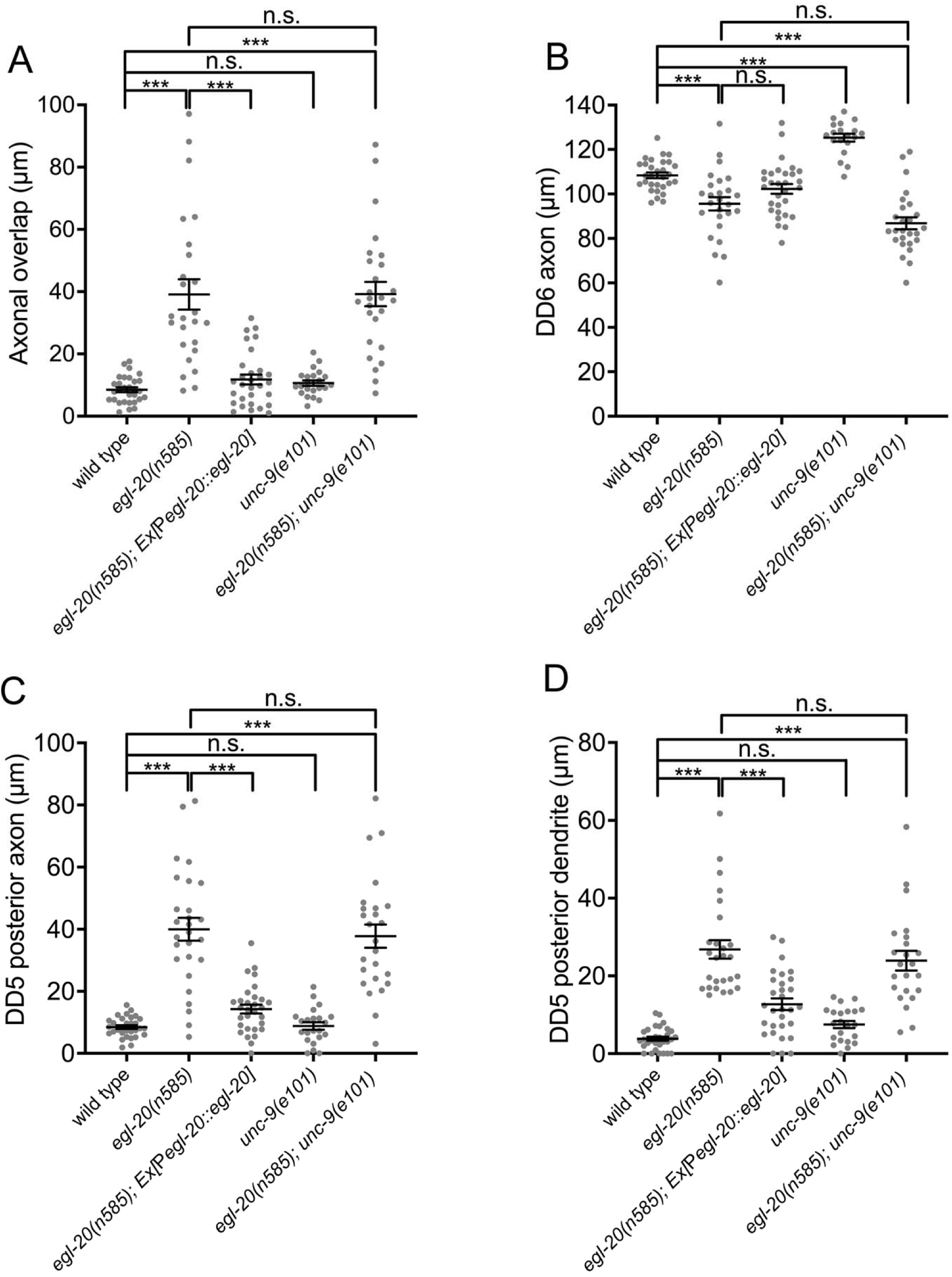
Expression of *egl-20* from its endogenous promoter rescues axonal tiling defects of *egl-20(n585)*. (A) Quantification of axonal overlap between DD5 and DD6. (B) Quantification of DD6 axonal length. (C) Quantification of DD5 posterior axonal length. (D) Quantification of DD5 posterior dendrite length. Each dot represents a single animal. Black bars indicate mean ± SEM. n.s.: not significant; ***p<0.001.

**Supplemental figure 2-1:**
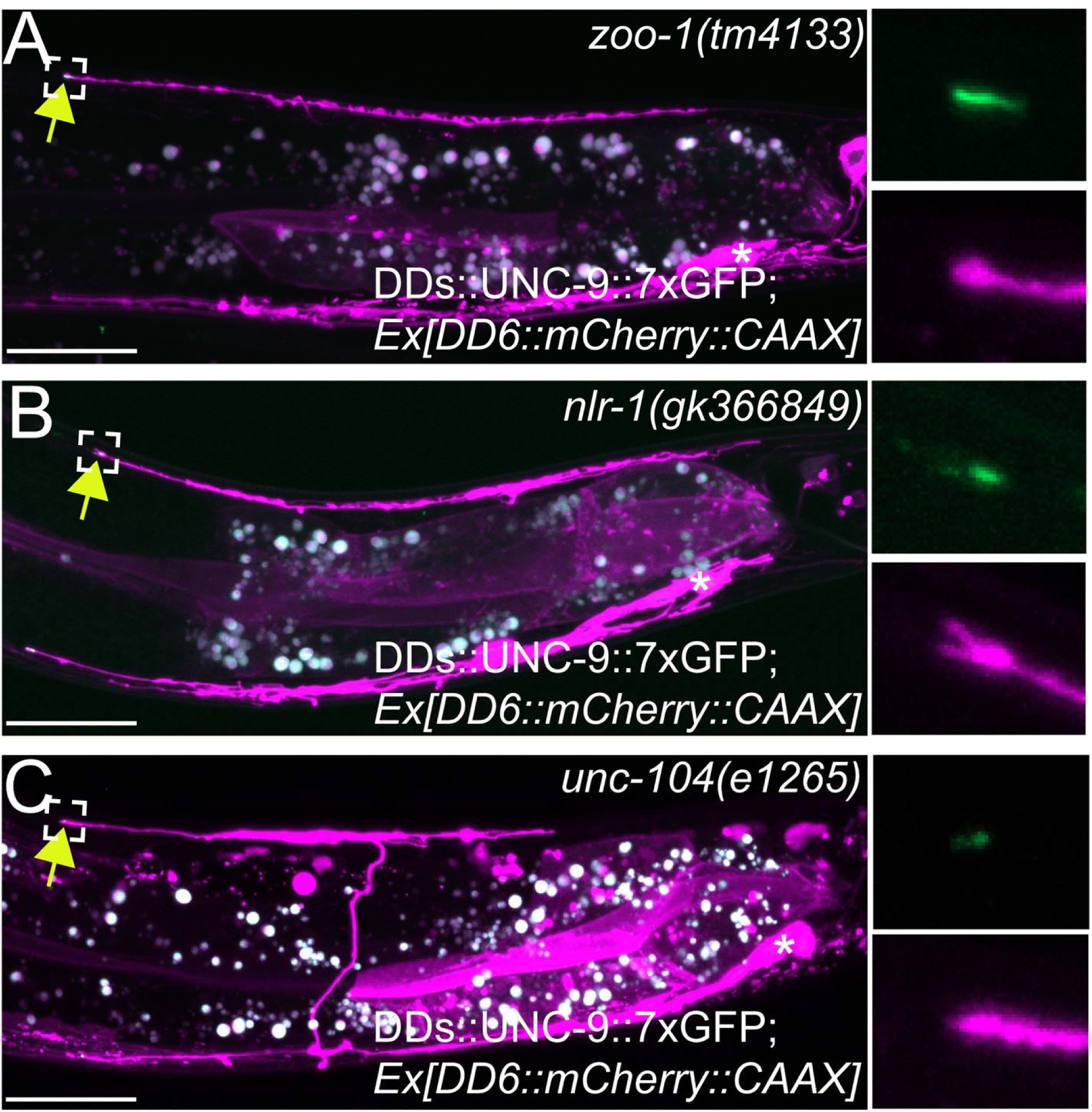
UNC-9 localization is not affected in *zoo-1(tm4133), nlr-1(gk366849),* and *unc-104(e1265)* mutants. (A-B) Representative image of UNC-9::7×GFP localization at the tip of the anterior DD6 axon in *zoo-1(tm4133)* (A), *nlr-1(gk366849)* (B), and *unc-104(e1265)* mutants. The magnified UNC-9::7×GFP and mCherry::CAAX signals, represented by the dotted box, at the anterior axon of DD6 are shown to the right of merged image. Asterisks: DD6 cell bodies. Scale bar: 20µm.

**Supplemental figures 2-2:**
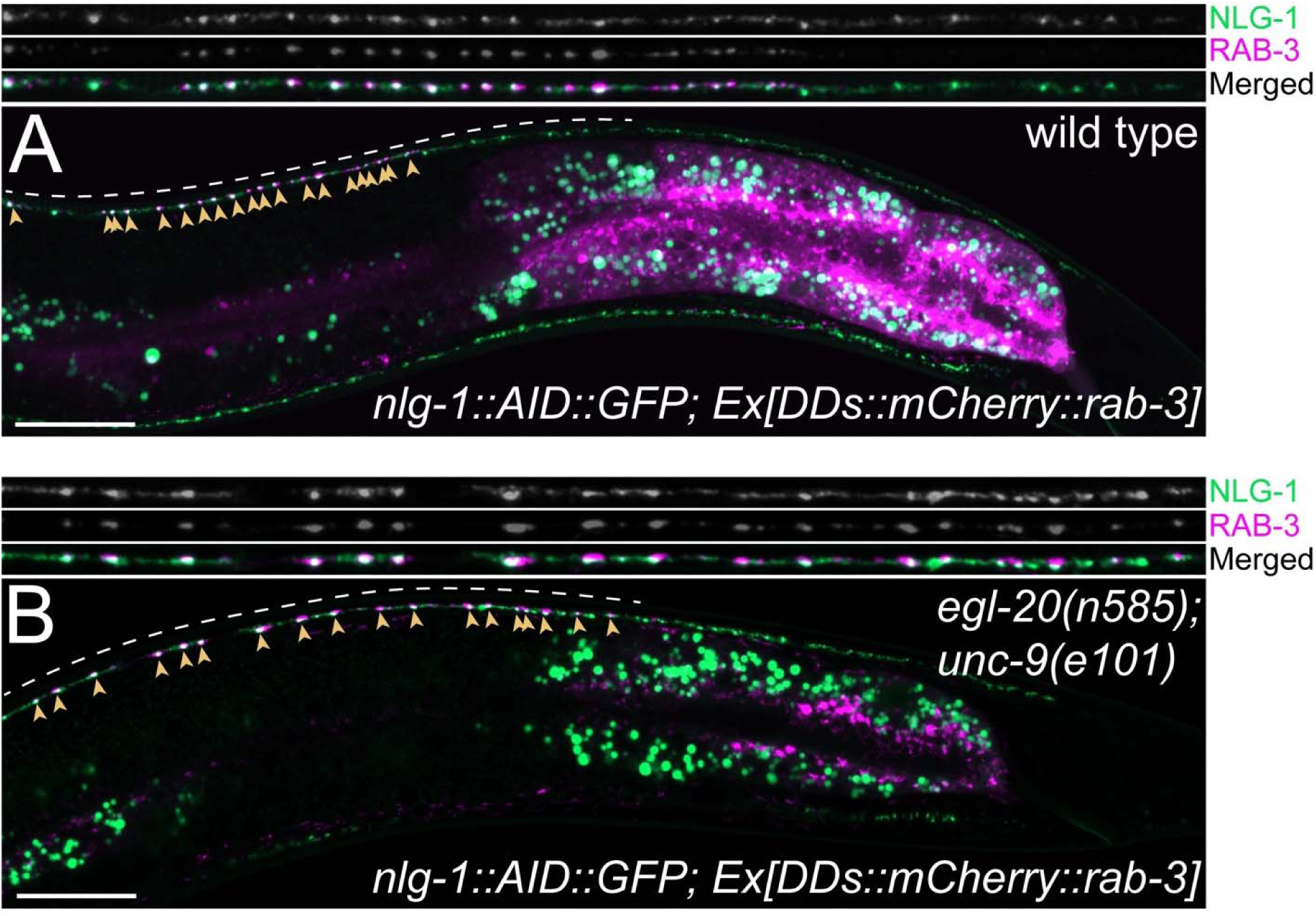
Co-localization between RAB-3 puncta in the DD5 posterior axon and the postsynaptic NLG-1. (A-B) Representative images of mCherry::RAB-3 expressed in DD5 and endogenous NLG-1::AID::GFP in wild type (A) and *egl-20(n585); unc-9(e101)* double mutants (B) . Arrowheads indicate mCherry::RAB-3 puncta. Straightened images of NLG-1::AID::GFP (first panel), mCherry::RAB-3 (second panel) and merged channels (third panel). Dotted lines indicate the straightened and magnified region of the dorsal nerve cord. Scale bar: 20µm.

**Supplemental figure 2-3:**
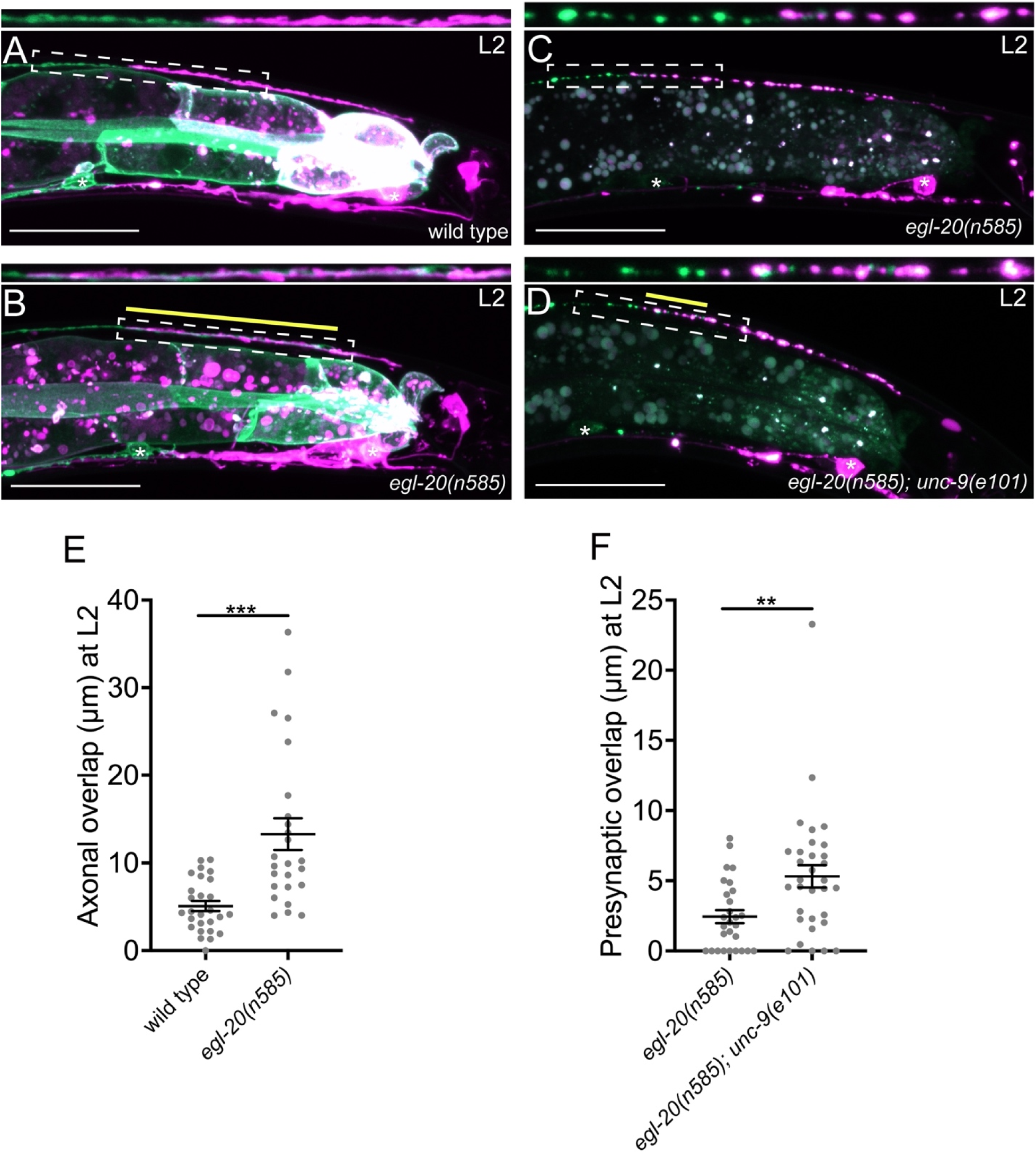
*egl-20* and *unc-9* are required for establishment of axonal and presynaptic tiling. (A-B) Representative image of axonal tiling in wild type (A) and *egl-20(n585)* (B) animals at the L2 stage. Yellow line represents region of axonal overlap between DD5 and DD6. (C-D) Representative image of presynaptic tiling in the *egl-20(n585)* (C), and *egl-20(n585); unc-9(e101)* (D) mutants at the L2 stage. Scale bar: 20µm. (E) Quantification of axonal overlap between DD5 and DD6. (F) Quantification of presynaptic overlap between DD5 and DD6. Asterisks: DD5 and DD6 cell bodies. Scale bar: 20µm. Each dot represents a single animal. Black bars indicate mean ± SEM. **p<0.01; ***p<0.001.

**Supplemental figure 3:**
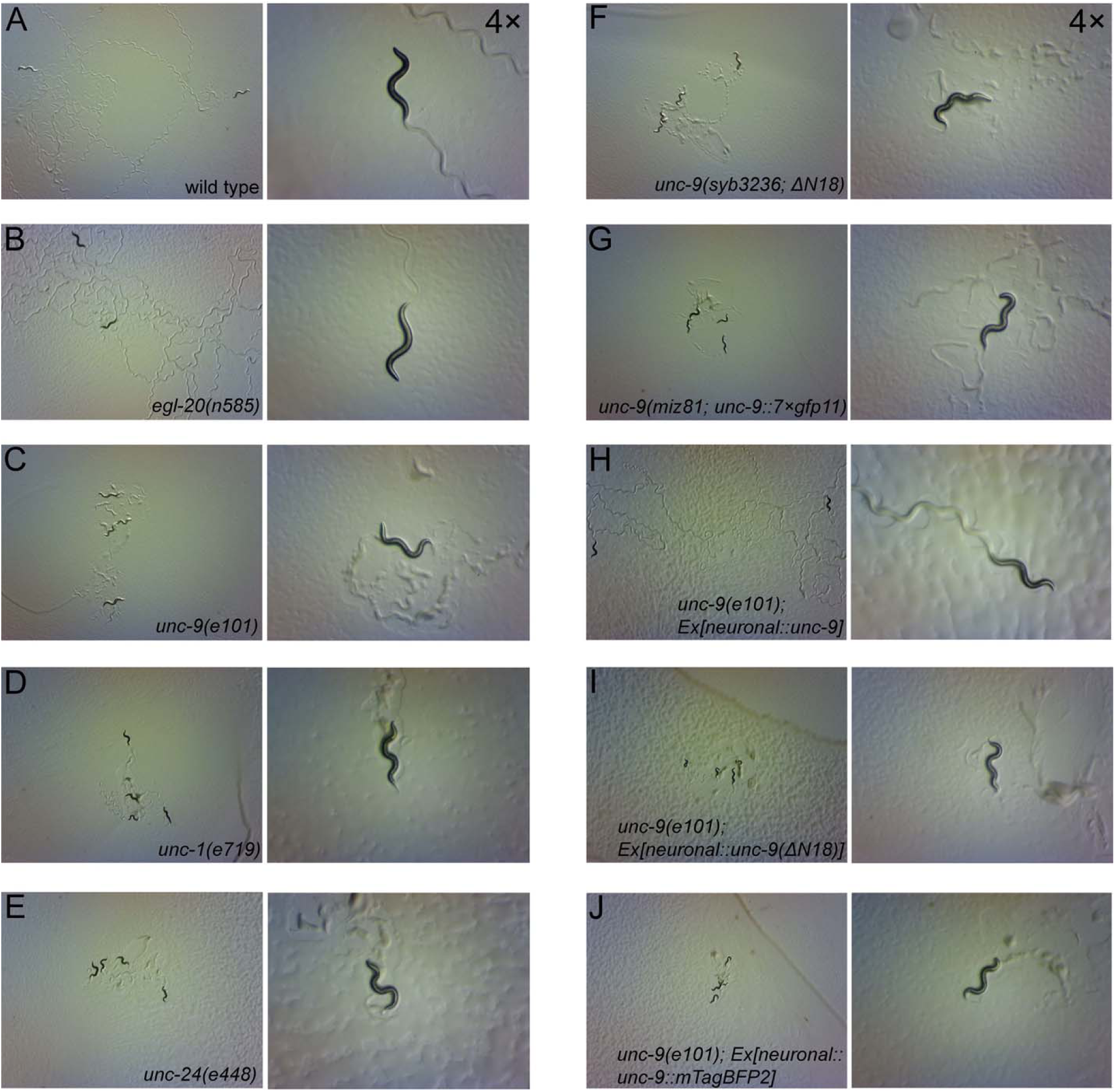
Locomotion defects of UNC-9-gap junction channel defective mutants. (A-J) Representative stereoscope images of wild type (A), *egl-20(n585)* (B), *unc-9(e101)* (C), *unc-1(e719)* (D), *unc-24(e448)* (E), *unc-9(syb3236; ΔN18)* (F), *unc-9(miz81; unc-9::7×gfp11)* (G), *unc-9(e101) Ex[neuronal::unc-9]* (H), *unc-9(e101) Ex[neuronal::unc-9(ΔN18)]* (I), *unc-9(e101) Ex[neuronal::unc-9::mTagBFP2]* (J)

## Notes

### Competing Interest Statement

The authors have declared no competing interest.

## References

Aiken, J., and Holzbaur, E.L.F. (2021). Cytoskeletal regulation guides neuronal trafficking to effectively supply the synapse. Curr Biol 31, R633–R650.

Altun, Z.F., Chen, B., Wang, Z.W., and Hall, D.H. (2009). High resolution map of Caenorhabditis elegans gap junction proteins. Dev Dyn 238, 1936–1950.

Au, V., Li-Leger, E., Raymant, G., Flibotte, S., Chen, G., Martin, K., Fernando, L., Doell, C., Rosell, F.I., Wang, S., et al. (2019). CRISPR/Cas9 Methodology for the Generation of Knockout Deletions in Caenorhabditis elegans. G3 (Bethesda) 9, 135–144.

Barbagallo, B., Philbrook, A., Touroutine, D., Banerjee, N., Oliver, D., Lambert, C.M., and Francis, M.M. (2017). Excitatory neurons sculpt GABAergic neuronal connectivity in the C. elegans motor circuit. Development 144, 1807–1819.

Brenner, S. (1974). The genetics of Caenorhabditis elegans. Genetics 77, 71–94.

Burendei, B., Shinozaki, R., Watanabe, M., Terada, T., Tani, K., Fujiyoshi, Y., and Oshima, A. (2020). Cryo-EM structures of undocked innexin-6 hemichannels in phospholipids. Sci Adv 6, eaax3157.

Cameron, S., and Rao, Y. (2010). Molecular mechanisms of tiling and self-avoidance in neural development. Mol Brain 3, 28.

Chen, B., Liu, Q., Ge, Q., Xie, J., and Wang, Z.W. (2007). UNC-1 regulates gap junctions important to locomotion in C. elegans. Curr Biol 17, 1334–1339.

Chen, X., Shibata, A.C., Hendi, A., Kurashina, M., Fortes, E., Weilinger, N.L., MacVicar, B.A., Murakoshi, H., and Mizumoto, K. (2018). Rap2 and TNIK control Plexin-dependent tiled synaptic innervation in C. elegans. Elife 7.

Dbouk, H.A., Mroue, R.M., El-Sabban, M.E., and Talhouk, R.S. (2009). Connexins: a myriad of functions extending beyond assembly of gap junction channels. Cell Commun Signal 7, 4.

Deng, Z., He, Z., Maksaev, G., Bitter, R.M., Rau, M., Fitzpatrick, J.A.J., and Yuan, P. (2020). Cryo-EM structures of the ATP release channel pannexin 1. Nat Struct Mol Biol 27, 373–381.

Dokshin, G.A., Ghanta, K.S., Piscopo, K.M., and Mello, C.C. (2018). Robust Genome Editing with Short Single-Stranded and Long, Partially Single-Stranded DNA Donors in Caenorhabditis elegans. Genetics 210, 781–787.

Elias, L.A., Wang, D.D., and Kriegstein, A.R. (2007). Gap junction adhesion is necessary for radial migration in the neocortex. Nature 448, 901–907.

Emoto, K., He, Y., Ye, B., Grueber, W.B., Adler, P.N., Jan, L.Y., and Jan, Y.N. (2004). Control of dendritic branching and tiling by the Tricornered-kinase/Furry signaling pathway in Drosophila sensory neurons. Cell 119, 245–256.

Emoto, K., Parrish, J.Z., Jan, L.Y., and Jan, Y.N. (2006). The tumour suppressor Hippo acts with the NDR kinases in dendritic tiling and maintenance. Nature 443, 210–213.

Falk, L., Dang-Lawson, M., Vega, J.L., Pournia, F., Choi, K., Jang, C., Naus, C.C., and Matsuuchi, L. (2014). Mutations of Cx43 that affect B cell spreading in response to BCR signaling. Biol Open 3, 185–194.

Fatemi, S.H., Folsom, T.D., Reutiman, T.J., and Lee, S. (2008). Expression of astrocytic markers aquaporin 4 and connexin 43 is altered in brains of subjects with autism. Synapse 62, 501–507.

Fire, A. (1986). Integrative transformation of Caenorhabditis elegans. EMBO J 5, 2673–2680.

Friedland, A.E., Tzur, Y.B., Esvelt, K.M., Colaiacovo, M.P., Church, G.M., and Calarco, J.A. (2013). Heritable genome editing in C. elegans via a CRISPR-Cas9 system. Nat Methods 10, 741–743.

Fuerst, P.G., Bruce, F., Tian, M., Wei, W., Elstrott, J., Feller, M.B., Erskine, L., Singer, J.H., and Burgess, R.W. (2009). DSCAM and DSCAML1 function in self-avoidance in multiple cell types in the developing mouse retina. Neuron 64, 484–497.

Fuerst, P.G., Koizumi, A., Masland, R.H., and Burgess, R.W. (2008). Neurite arborization and mosaic spacing in the mouse retina require DSCAM. Nature 451, 470–474.

Giaume, C., Saez, J.C., Song, W., Leybaert, L., and Naus, C.C. (2019). Connexins and pannexins in Alzheimer’s disease. Neurosci Lett 695, 100–105.

Gibson, D.G. (2011). Enzymatic assembly of overlapping DNA fragments. Methods Enzymol 498, 349–361.

Grueber, W.B., Jan, L.Y., and Jan, Y.N. (2002). Tiling of the Drosophila epidermis by multidendritic sensory neurons. Development 129, 2867–2878.

Grueber, W.B., and Sagasti, A. (2010). Self-avoidance and tiling: Mechanisms of dendrite and axon spacing. Cold Spring Harb Perspect Biol 2, a001750.

Grueber, W.B., Ye, B., Moore, A.W., Jan, L.Y., and Jan, Y.N. (2003). Dendrites of distinct classes of Drosophila sensory neurons show different capacities for homotypic repulsion. Curr Biol 13, 618–626.

Guilmatre, A., Dubourg, C., Mosca, A.L., Legallic, S., Goldenberg, A., Drouin-Garraud, V., Layet, V., Rosier, A., Briault, S., Bonnet-Brilhault, F., et al. (2009). Recurrent rearrangements in synaptic and neurodevelopmental genes and shared biologic pathways in schizophrenia, autism, and mental retardation. Arch Gen Psychiatry 66, 947–956.

Hall, D.H. (2017). Gap junctions in C. elegans: Their roles in behavior and development. Dev Neurobiol 77, 587–596.

Hall, D.H., and Hedgecock, E.M. (1991). Kinesin-related gene unc-104 is required for axonal transport of synaptic vesicles in C. elegans. Cell 65, 837–847.

He, S., Cuentas-Condori, A., and Miller, D.M., 3rd (2019). NATF (Native and Tissue-Specific Fluorescence): A Strategy for Bright, Tissue-Specific GFP Labeling of Native Proteins in Caenorhabditis elegans. Genetics 212, 387–395.

Hendi, A., Kurashina, M., and Mizumoto, K. (2019). Intrinsic and extrinsic mechanisms of synapse formation and specificity in C. elegans. Cell Mol Life Sci 76, 2719–2738.

Inaki, M., Yoshikawa, S., Thomas, J.B., Aburatani, H., and Nose, A. (2007). Wnt4 is a local repulsive cue that determines synaptic target specificity. Curr Biol 17, 1574–1579.

Jang, H., Levy, S., Flavell, S.W., Mende, F., Latham, R., Zimmer, M., and Bargmann, C.I. (2017). Dissection of neuronal gap junction circuits that regulate social behavior in Caenorhabditis elegans. Proc Natl Acad Sci U S A 114, E1263–E1272.

Kameritsch, P., Khandoga, N., Pohl, U., and Pogoda, K. (2013). Gap junctional communication promotes apoptosis in a connexin-type-dependent manner. Cell Death Dis 4, e584.

Kameritsch, P., Pogoda, K., and Pohl, U. (2012). Channel-independent influence of connexin 43 on cell migration. Biochim Biophys Acta 1818, 1993–2001.

Kawano, T., Po, M.D., Gao, S., Leung, G., Ryu, W.S., and Zhen, M. (2011). An imbalancing act: gap junctions reduce the backward motor circuit activity to bias C. elegans for forward locomotion. Neuron 72, 572–586.

Klassen, M.P., and Shen, K. (2007). Wnt signaling positions neuromuscular connectivity by inhibiting synapse formation in C. elegans. Cell 130, 704–716.

Lapato, A.S., and Tiwari-Woodruff, S.K. (2018). Connexins and pannexins: At the junction of neuro-glial homeostasis & disease. J Neurosci Res 96, 31–44.

Lasseigne, A.M., Echeverry, F.A., Ijaz, S., Michel, J.C., Martin, E.A., Marsh, A.J., Trujillo, E., Marsden, K.C., Pereda, A.E., and Miller, A.C. (2021). Electrical synaptic transmission requires a postsynaptic scaffolding protein. Elife 10.

Liu, P., Chen, B., and Wang, Z.W. (2011). Gap junctions synchronize action potentials and Ca2+ transients in Caenorhabditis elegans body wall muscle. J Biol Chem 286, 44285–44293.

Liu, Q., Chen, B., Gaier, E., Joshi, J., and Wang, Z.W. (2006). Low conductance gap junctions mediate specific electrical coupling in body-wall muscle cells of Caenorhabditis elegans. J Biol Chem 281, 7881–7889.

Machtaler, S., Dang-Lawson, M., Choi, K., Jang, C., Naus, C.C., and Matsuuchi, L. (2011). The gap junction protein Cx43 regulates B-lymphocyte spreading and adhesion. J Cell Sci 124, 2611–2621.

Maro, G.S., Gao, S., Olechwier, A.M., Hung, W.L., Liu, M., Ozkan, E., Zhen, M., and Shen, K. (2015). MADD-4/Punctin and Neurexin Organize C. elegans GABAergic Postsynapses through Neuroligin. Neuron 86, 1420–1432.

Maro, G.S., Klassen, M.P., and Shen, K. (2009). A beta-catenin-dependent Wnt pathway mediates anteroposterior axon guidance in C. elegans motor neurons. PLoS One 4, e4690.

McDiarmid, T.A., Belmadani, M., Liang, J., Meili, F., Mathews, E.A., Mullen, G.P., Hendi, A., Wong, W.R., Rand, J.B., Mizumoto, K., et al. (2020). Systematic phenomics analysis of autism-associated genes reveals parallel networks underlying reversible impairments in habituation. Proc Natl Acad Sci U S A 117, 656–667.

Mello, C.C., Kramer, J.M., Stinchcomb, D., and Ambros, V. (1991). Efficient gene transfer in C.elegans: extrachromosomal maintenance and integration of transforming sequences. EMBO J 10, 3959–3970.

Meng, L., and Yan, D. (2020). NLR-1/CASPR Anchors F-Actin to Promote Gap Junction Formation. Dev Cell 55, 574–587 e573.

Miao, G., Godt, D., and Montell, D.J. (2020). Integration of Migratory Cells into a New Site In Vivo Requires Channel-Independent Functions of Innexins on Microtubules. Dev Cell 54, 501–515 e509.

Michalski, K., Syrjanen, J.L., Henze, E., Kumpf, J., Furukawa, H., and Kawate, T. (2020). The Cryo-EM structure of pannexin 1 reveals unique motifs for ion selection and inhibition. Elife 9.

Millard, S.S., Flanagan, J.J., Pappu, K.S., Wu, W., and Zipursky, S.L. (2007). Dscam2 mediates axonal tiling in the Drosophila visual system. Nature 447, 720–724.

Mitchell, K.J. (2011). The genetics of neurodevelopmental disease. Curr Opin Neurobiol 21, 197–203.

Mizumoto, K., and Shen, K. (2013a). Interaxonal interaction defines tiled presynaptic innervation in C. elegans. Neuron 77, 655–666.

Mizumoto, K., and Shen, K. (2013b). Two Wnts instruct topographic synaptic innervation in C. elegans. Cell Rep 5, 389–396.

Mok, D.Z., Sternberg, P.W., and Inoue, T. (2015). Morphologically defined sub-stages of C. elegans vulval development in the fourth larval stage. BMC Dev Biol 15, 26.

Norris, A.D., Kim, H.M., Colaiacovo, M.P., and Calarco, J.A. (2015). Efficient Genome Editing in Caenorhabditis elegans with a Toolkit of Dual-Marker Selection Cassettes. Genetics 201, 449–458.

Obinata, H., Sugimoto, A., and Niwa, S. (2018). Streptothricin acetyl transferase 2 (Sat2): A dominant selection marker for Caenorhabditis elegans genome editing. PLoS One 13, e0197128.

Olk, S., Zoidl, G., and Dermietzel, R. (2009). Connexins, cell motility, and the cytoskeleton. Cell Motil Cytoskeleton 66, 1000–1016.

Onishi, K., Hollis, E., and Zou, Y. (2014). Axon guidance and injury-lessons from Wnts and Wnt signaling. Curr Opin Neurobiol 27, 232–240.

Oshima, A., Matsuzawa, T., Murata, K., Tani, K., and Fujiyoshi, Y. (2016a). Hexadecameric structure of an invertebrate gap junction channel. J Mol Biol 428, 1227–1236.

Oshima, A., Tani, K., and Fujiyoshi, Y. (2016b). Atomic structure of the innexin-6 gap junction channel determined by cryo-EM. Nat Commun 7, 13681.

Palumbos, S.D., Skelton, R., McWhirter, R., Mitchell, A., Swann, I., Heifner, S., Von Stetina, S., and Miller, D.M., 3rd (2021). cAMP controls a trafficking mechanism that maintains the neuron specificity and subcellular placement of electrical synapses. Dev Cell 56, 3235–3249 e3234.

Pereda, A.E. (2014). Electrical synapses and their functional interactions with chemical synapses. Nat Rev Neurosci 15, 250–263.

Philbrook, A., Ramachandran, S., Lambert, C.M., Oliver, D., Florman, J., Alkema, M.J., Lemons, M., and Francis, M.M. (2018). Neurexin directs partner-specific synaptic connectivity in C. elegans. Elife 7.

Poon, V.Y., Klassen, M.P., and Shen, K. (2008). UNC-6/netrin and its receptor UNC-5 locally exclude presynaptic components from dendrites. Nature 455, 669–673.

Sanchez, A., Castro, C., Flores, D.L., Gutierrez, E., and Baldi, P. (2019). Gap Junction Channels of Innexins and Connexins: Relations and Computational Perspectives. Int J Mol Sci 20.

Sanchez-Arias, J.C., Candlish, R.C., van der Slagt, E., and Swayne, L.A. (2020). Pannexin 1 Regulates Dendritic Protrusion Dynamics in Immature Cortical Neurons. eNeuro 7.

Sanchez-Arias, J.C., Liu, M., Choi, C.S.W., Ebert, S.N., Brown, C.E., and Swayne, L.A. (2019). Pannexin 1 Regulates Network Ensembles and Dendritic Spine Development in Cortical Neurons. eNeuro 6.

Sanes, J.R., and Yamagata, M. (2009). Many paths to synaptic specificity. Annu Rev Cell Dev Biol 25, 161–195.

Shao, Q., Lindstrom, K., Shi, R., Kelly, J., Schroeder, A., Juusola, J., Levine, K.L., Esseltine, J.L., Penuela, S., Jackson, M.F., et al. (2016). A Germline Variant in the PANX1 Gene Has Reduced Channel Function and Is Associated with Multisystem Dysfunction. J Biol Chem 291, 12432–12443.

Sudhof, T.C. (2008). Neuroligins and neurexins link synaptic function to cognitive disease. Nature 455, 903–911.

Swayne, L.A., and Bennett, S.A. (2016). Connexins and pannexins in neuronal development and adult neurogenesis. BMC Cell Biol 17 *Suppl 1*, 10.

Tabuchi, K., Blundell, J., Etherton, M.R., Hammer, R.E., Liu, X., Powell, C.M., and Sudhof, T.C. (2007). A neuroligin-3 mutation implicated in autism increases inhibitory synaptic transmission in mice. Science (New York, NY) 318, 71–76.

Tang, G., Gudsnuk, K., Kuo, S.H., Cotrina, M.L., Rosoklija, G., Sosunov, A., Sonders, M.S., Kanter, E., Castagna, C., Yamamoto, A., et al. (2014). Loss of mTOR-dependent macroautophagy causes autistic-like synaptic pruning deficits. Neuron 83, 1131–1143.

Ting, C.Y., Herman, T., Yonekura, S., Gao, S., Wang, J., Serpe, M., O’Connor, M.B., Zipursky, S.L., and Lee, C.H. (2007). Tiling of r7 axons in the Drosophila visual system is mediated both by transduction of an activin signal to the nucleus and by mutual repulsion. Neuron 56, 793–806.

Tran, T.S., Rubio, M.E., Clem, R.L., Johnson, D., Case, L., Tessier-Lavigne, M., Huganir, R.L., Ginty, D.D., and Kolodkin, A.L. (2009). Secreted semaphorins control spine distribution and morphogenesis in the postnatal CNS. Nature 462, 1065–1069.

Tu, H., Pinan-Lucarre, B., Ji, T., Jospin, M., and Bessereau, J.L. (2015). C. elegans Punctin Clusters GABA(A) Receptors via Neuroligin Binding and UNC-40/DCC Recruitment. Neuron 86, 1407–1419.

Wen, Z., Nguyen, H.N., Guo, Z., Lalli, M.A., Wang, X., Su, Y., Kim, N.S., Yoon, K.J., Shin, J., Zhang, C., et al. (2014). Synaptic dysregulation in a human iPS cell model of mental disorders. Nature 515, 414–418.

Whangbo, J., and Kenyon, C. (1999). A Wnt signaling system that specifies two patterns of cell migration in C. elegans. Mol Cell 4, 851–858.

White, J.G., Albertson, D.G., and Anness, M.A. (1978). Connectivity changes in a class of motoneurone during the development of a nematode. Nature 271, 764–766.

White, J.G., Southgate, E., Thomson, J.N., and Brenner, S. (1976). The structure of the ventral nerve cord of Caenorhabditis elegans. Philosophical transactions of the Royal Society of London Series B, Biological sciences 275, 327–348.

White, J.G., Southgate, E., Thomson, J.N., and Brenner, S. (1986). The structure of the nervous system of the nematode Caenorhabditis elegans. Philosophical transactions of the Royal Society of London Series B, Biological sciences 314, 1–340.

Yeh, E., Kawano, T., Ng, S., Fetter, R., Hung, W., Wang, Y., and Zhen, M. (2009). Caenorhabditis elegans innexins regulate active zone differentiation. J Neurosci 29, 5207–5217.

Yogev, S., and Shen, K. (2014). Cellular and molecular mechanisms of synaptic specificity. Annu Rev Cell Dev Biol 30, 417–437.

Zipursky, S.L., and Grueber, W.B. (2013). The molecular basis of self-avoidance. Annu Rev Neurosci 36, 547–568.

Zou, Y. (2004). Wnt signaling in axon guidance. Trends Neurosci 27, 528–532.

